# Discovery of a novel UV-absorbing mycosporine-like amino acid in *Vertebrata lanosa* using an expanded combinatorial structure database incorporating non-proteinogenic amino acids and organic solutes

**DOI:** 10.64898/2026.08.25.746913

**Authors:** Armin Oberosler, Fabian Jürgen Hammerle, Sebastian Lanner, Hossam Elgabarty, Solène Connan, Fernanda Pita, Barış Ballık, Ulf Karsten, Markus Ganzera

## Abstract

Mycosporine-like amino acids (MAAs) are among nature’s most effective sunscreen compounds, capable of converting harmful ultraviolet radiation into harmless heat, and are widely distributed in marine organisms such as red macroalgae. Although decades of research have led to numerous discoveries, the rate of new MAA identifications has declined. To address this, we considerably expanded our previously developed combinatorial MAA database, increasing the number of covered structures tenfold. Following a comprehensive literature search for plausible but undescribed building blocks, the database now incorporates an extensive set of proteinogenic and non-proteinogenic amino acids, as well as other marine organic osmolytes, in combination with all (currently) known MAA scaffolds. This expanded resource was integrated into our identification platform, which combines UHPLC-VWD-HRMS^2^ analysis, feature-based molecular networking, and bioinformatics-driven annotation. Application of this updated workflow enabled the isolation and structural elucidation of a novel MAA, mycosporine-cysteinolic acid, from the red marine macroalga *Vertebrata lanosa*. Altogether, this study provides a valuable extension of the bioinformatics-based MAA screening pipeline, enhancing the annotation and discovery of novel MAAs in natural matrices.

**Graphical abstract:** 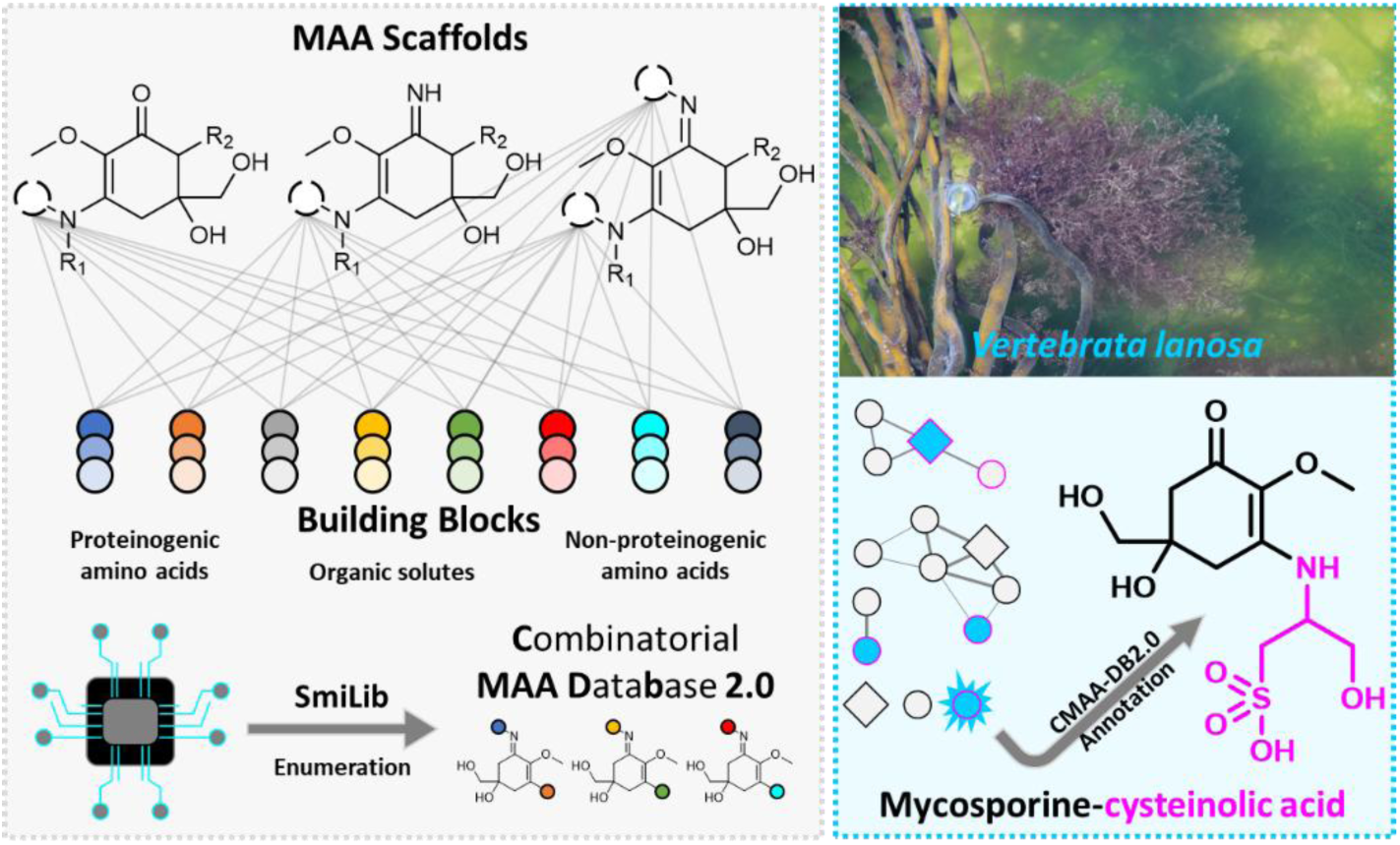

## 1 Introduction

Solar ultraviolet (UV) radiation is a major environmental stressor for photosynthetic organisms, particularly in shallow water and intertidal marine habitats where exposure to high irradiance and periodic desiccation can intensify photodamage. Excessive UV-A and UV-B radiation can impair photosynthesis, damage cellular macromolecules, and promote oxidative stress through the formation of reactive oxygen species [1, 2]. Marine algae have therefore evolved multiple photoprotective mechanisms, including antioxidant systems, repair mechanisms, and the accumulation of UV-absorbing secondary metabolites [3, 4]. Among these natural sunscreens, mycosporine-like amino acids (MAAs) are especially relevant. They are low-molecular-weight, hydrophilic compounds that are widely distributed throughout marine food webs, from primary producers to fish [5, 6]. They have attracted considerable attention due to their diverse bioactivities, including antioxidant and wound-healing properties [7, 8], but are best known for their ability to absorb harmful UV radiation [9]. Most MAAs exhibit absorption maxima between 310 and 360 nm, possess exceptionally high molar extinction coefficients, are highly photostable, and efficiently dissipate absorbed energy as heat, making them one of the most widespread natural sunscreen systems in aquatic organisms [10–12]. Consequently, MAAs have gained increasing interest in the cosmetic industry as environmentally friendly and biocompatible UV filters. Commercial formulations such as HELIONORI^®^ (Gelyma, France) and Helioguard^®^ 365 (Mibelle Group, Switzerland), based on the MAAs shinorine and porphyra-334 present in the red macroalga *Porphyra umbilicalis*, exemplify their applied potential [13, 14]. Overall, the combination of diverse biological activities and excellent biocompatibility makes them promising multifunctional compounds for cosmetic, pharmaceutical, and biotechnological applications [15, 16]. Despite their promising properties, MAAs have not yet achieved broad commercial utilization. This is partly due to limitations in their physicochemical properties, particularly their high hydrophilicity, which complicates their purification from extracts and formulation, as well as the lack of viable strategies for their *de novo* chemical synthesis. To date, only White et al. have described the synthesis of mycosporin I and mycosporine-glycine from (D)-(-)-quinic acid via an iminophosphorane intermediate, involving 14 steps and resulting in an overall yield of less than 1% [17]. As a result, MAAs must still be tediously isolated from natural sources or used in the form of algal extracts.

Despite decades of research, the discovery of novel MAAs remains a persistent challenge in natural product chemistry. To date, more than 70 distinct MAAs have been identified [15], yet the discovery of new structures has stagnated in recent years. Early approaches have often relied on random success, which, while occasionally fruitful, are unpredictable and inefficient. More directed strategies focused on exploring organisms closely related to species known to produce various MAAs, leveraging evolutionary and biochemical similarities to increase the likelihood of identifying new compounds. The observed stagnation can largely be attributed to two factors: easily available biomaterials have already been thoroughly characterized, while less accessible organisms often yield insufficient quantities of MAAs for isolation and structure elucidation. In addition, some MAAs occur at low concentrations, and their abundance can vary significantly depending on environmental conditions, developmental stage, and extraction protocols [18–20]. Traditional workflows for the isolation and structural elucidation of MAAs are well established and generally involve extraction of dried biomaterial with polar solvents, followed by stepwise purification using ion-exchange techniques and chromatographic methods such as medium-pressure liquid chromatography (MPLC) or semi-preparative high-performance liquid chromatography (HPLC) [21, 22]. Additional purification steps, including size-exclusion chromatography or hydrophilic interaction liquid chromatography (HILIC), may be required for certain compounds [23]. While such approaches have successfully led to the identification of novel MAAs in recent years, for example, in the red algal genus *Bostrychia* sp., which typically occurs as epiphyte on mangrove trees [24, 25], in the green algal genus *Klebsormidium* sp., mainly exhibiting a terrestrial lifestyle in biological soil crusts [26], and in the recently described green alga *Apatococcus ammoniophilus*, that grows as terrestrial epiphyte on temperate pine and other trees [23], the overall rate of discovery has dropped. All of the mentioned algal taxa prefer aeroterrestrial habitats, and hence are regularly exposed to high insolation and desiccation.

To tackle this decline in the discovery of MAAs, metabolomics-based approaches have recently been introduced into the field of algal research. In particular, feature-based molecular networking (FBMN) combined with *in silico* annotation tools enables the detection of structurally related compounds even at low abundance [20, 27]. Building on this, we previously developed a combinatorial structure database of theoretically possible MAAs, constructed from known core structures and a small set of proteinogenic and selected non-proteinogenic amino acids [23]. This database enables the generation of plausible structural candidates following FBMN analysis, thereby increasing annotation confidence and supporting subsequent structural characterization. The application of this informed strategy to extracts of the basidiomycete *Stereum gausapatum* and the microalga *A. ammoniophilus* has proven successful, leading to the isolation and identification of the major MAAs mycosporine-serinol and algasporine-glycine, and thus representing an important step toward a more systematic identification platform. In addition, our extensive studies on MAA profiles have established robust chromatographic purification strategies that facilitate their isolation [22, 24, 28–31].

In the present study, we expand this concept by substantially broadening the chemical search space of the combinatorial MAA database through the incorporation of additional naturally occurring nitrogen-containing metabolites, thereby better reflecting the metabolic diversity of marine organisms, which rely on a finely tuned network of metabolites to cope with environmental stressors such as osmotic stress and UV radiation. Osmotic stress is mitigated by organic compatible solutes [32–34], including amino acids, betaines, and polyols, whereas UV protection is primarily mediated by MAAs [35, 36]. Since MAAs comprise a (deoxy)gadusol core conjugated to amino acids or other amine-containing compounds, we hypothesized that compatible solutes with primary or secondary amino groups may act as alternative building blocks for previously unrecognized MAA structures.

To test this hypothesis, we confirmed the presence of the MAA substituents serinol, *N*-methylglycine, and *N*-methylserine in extracts of *S. gausapatum*, *A. ammoniophilus*, and *Klebsormidium crenulatum*. Subsequently, we developed an updated combinatorial MAA database (CMAA-DB2.0) and integrated it into a metabolomics workflow combining UHPLC-VWD-HRMS^2^ analysis, feature-based molecular networking (FBMN), and annotation using the SIRIUS software platform. As a proof of concept, this approach was applied to a polar extract of the marine red macroalga *Vertebrata lanosa*, an obligate epiphyte on the brown seaweed *Ascophyllum nodosum*, enabling the detection, isolation, and structural elucidation of a novel MAA. Here, we describe the rationale and construction of the updated CMAA-DB 2.0, the metabolomic and chemical investigation of *V. lanosa*, the isolation of several known MAAs alongside the novel compound mycosporine-cysteinolic acid, including the elucidation of its structure and absolute configuration, and its distribution across different algal extracts.

## 2 Results and Discussion

MAAs are biosynthesized via the shikimate and/or pentose phosphate pathways [37–39]; however, the exact route remains incompletely understood, particularly with respect to recently discovered and structurally unusual compounds such as klebsormidin A and B and algasporine-glycine, which feature a gadusol core with *N*-methylated substituents. One unresolved question is whether *N*-methylation occurs after condensation with the amino acid or whether *N*-methylated amino acids are directly incorporated during biosynthesis. To preliminarily address this question, we revisited metabolomic datasets of *Stereum gausapatum*, *Apatococcus ammoniophilus*, and *Klebsormidium crenulatum*, specifically screening for the presence of structural motifs corresponding to serinol, *N*-methylglycine (sarcosine), and *N*-methylserine.

Quasimolecular ions, i.e., [M+H]^+^ adducts, corresponding to the targeted substituents were detected within a mass window of 10 ppm, co-eluting with the respective injection peaks in the UHPLC runs (Figure 1). Signals consistent with serinol, *N*-methylglycine, and *N*-methylserine were observed. The ion tentatively assigned to serinol appeared at low intensity, precluding acquisition of a reliable MS^2^ spectrum. In contrast, ions putatively corresponding to *N*-methylglycine and *N*-methylserine were detected at higher intensities, enabling tandem MS analysis. The observed fragmentation patterns were consistent with publicly available reference spectra, suggesting that *N*-methylated amino acids are endogenously present and may be directly incorporated into MAAs during biosynthesis. In line with this, sarcosine has been reported to serve both as an osmolyte and a carbon source in the halophile bacterium *Vibrio natriegens* [40], indicating that such compounds can be readily available in marine organisms. Its occurrence and osmoregulatory role in *A. ammoniophilus* therefore appear plausible. Although rarely reported, *N*-methylserine has been identified as a component of cyclic heptapeptide hepatotoxins in the cyanobacterial genus *Nostoc* [41]. In general, amino acid methylation is a common biochemical process. The stepwise *N*-methylation of glycine during betaine biosynthesis in the diatom *Thalassiosira pseudonana* [42] supports the plausibility of such modifications in MAA biosynthesis.

**Figure 1.**
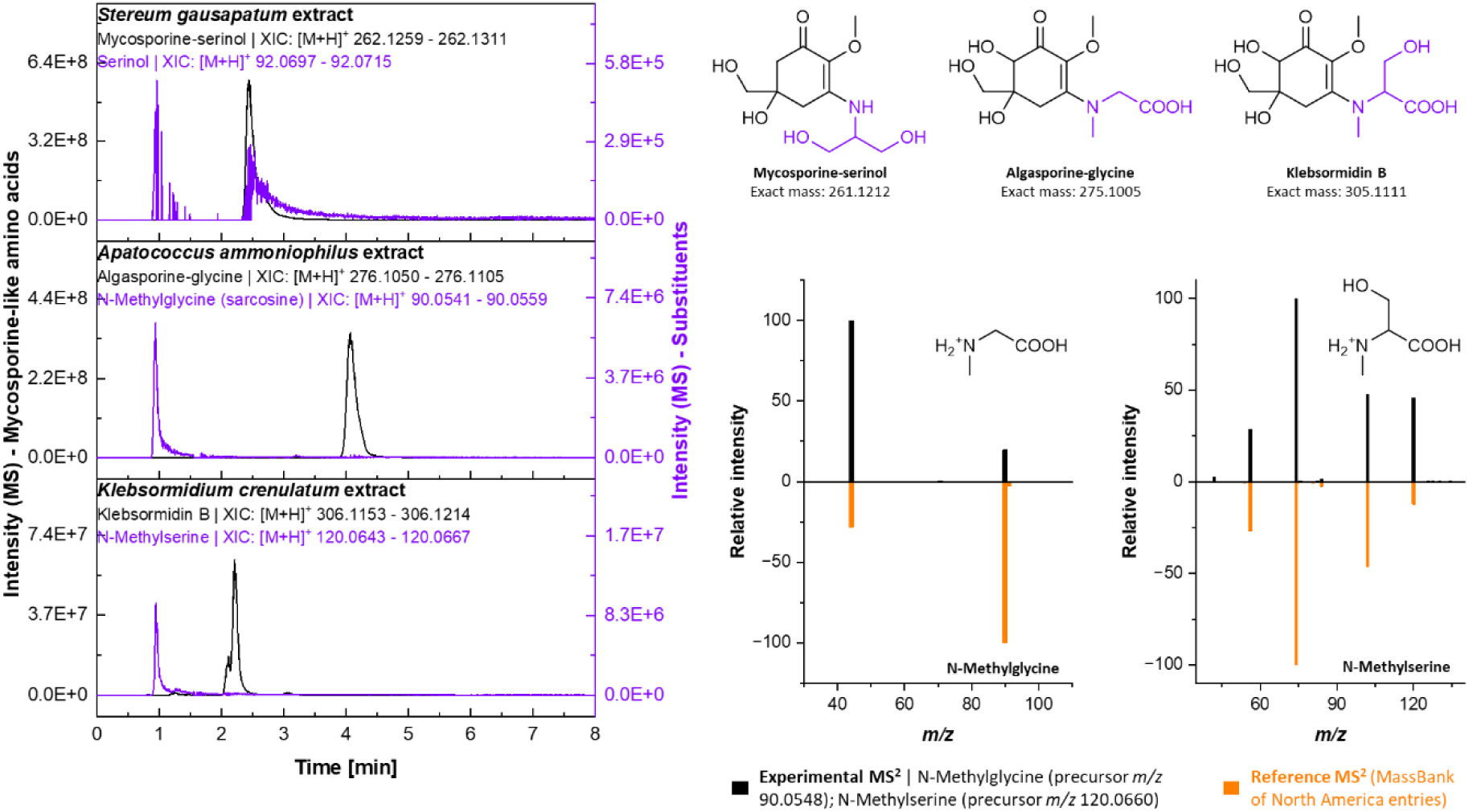
Extracted ion chromatograms (XICs) of quasimolecular ions ([M+H]^+^) corresponding to mycosporine-serinol, serinol, algasporine-glycine, *N*-methylglycine (sarcosine), *N*-methylserine, and klebsormidin B, generated using a 10 ppm mass window around the theoretical exact masses in extracts of *Stereum gausapatum*, *Apatococcus ammoniophilus*, and *Klebsormidium crenulatum*. Molecular structures, including exact masses, are shown for mycosporine-serinol, algasporine-glycine, and klebsormidin B. Experimental MS^2^ spectra of *N*-methylglycine and *N*-methylserine are compared with reference spectra from public databases (PubChem / MassBank of North America; *N*-methylglycine: splash10-0006-9000000000-067202e218085aadae08, *N*-methylserine: splash10-00di-9300000000-a496e99ad53b7b76d67f).

Motivated by both the structural diversity of known MAAs and this observation, we next considered additional potential substituents. Given that MAAs are characterized by amino acid-derived moieties and that many of them belong to the broader class of organic compatible solutes accumulated by marine organisms in response to osmotic stress, we systematically screened the literature for osmolytes and unusual non-proteinogenic amino acids bearing primary or secondary amino groups suitable for condensation with the (deoxy)gadusol core. For macroalgae, comprehensive overviews have been provided by Tarakhovskaya et al. and Harnedy & FitzGerald [43, 44]. Following the classification proposed by these authors, several compounds were considered as potential building blocks, including ureido amino acids and ornithine; β-alanine and its derivatives; proline-like secondary amino acids; kainoid amino acids; amino acid betaines; sulfur-containing amino acids; and iodo-amino acids. In addition, phosphorus-containing compounds such as phosphoserine and phosphoethanolamine, reported for example in *Porphyra* sp., were also added as possible candidates [45]. Another potential binding partner is cysteinolic acid, a sulfonate found in many marine and freshwater organisms. It was first isolated in 1957 from the red macroalga *Vertebrata lanosa* and described as an “internal salt” [46]. Fenizia and colleagues reported that cysteinolic acid levels in the diatom *Thalassiosira weissflogii* increased under elevated salinity, supporting its role in osmoadaptation [47]. To further expand the list of candidate building blocks, we also considered other extremophiles, including bacteria and archaea that inhabit environments similar to those of algae. Zwitterionic solutes such as (hydroxy)ectoine, *N*γ-acetyldiaminobutyrate, *N*ε-acetyl-β-lysine, and β-glutamine, which can be found in halotolerant or halophilic genera such as *Halomonas* and *Methanohalophilus*, emerged as promising candidates. Anionic solutes, including β-glutamate, which occurs in many halotolerant bacteria and methanogens, may also serve as MAA constituents [48]. A full list of considered metabolites is provided in the supporting information (Table S2). As outlined in our previous work, MAAs are characterized by a limited set of core scaffolds conjugated with various amine-containing substituents, predominantly amino acids, giving rise to their structural diversity. This modular architecture makes MAAs particularly well suited for the construction of combinatorial structure databases (Figure 2). Following the approach introduced by Nothias-Esposito et al. [49], who generated combinatorial libraries of theoretical premyrsinane and myrsinane derivatives, we constructed a set of SMILES input strings for the software SmiLib [50]. In the initial version of our combinatorial database, MAAs featuring hydrogen substitution (e.g., palythine) were not directly included and had to be added individually in a later step. To address this limitation, we expanded the list of core structures. Specifically, for the aminocyclohexenimine-, cyclohexenone-, and palythine-type cores, we incorporated *N*-methylhydroxy-, *N*-methyl-, and hydroxy-derivatives, resulting in four variations for each of the three core types (Figure 2). Additionally, several newly identified potential building blocks, such as proline, contain secondary amine functionalities that are often embedded within heterocyclic structures. To include these structures, the SmiLib input strategy had to be modified, as the software does not allow more than one atom to be attached to the spacer defining the linkage position on the scaffold (in this case, the nitrogen atom). Consequently, a separate set of SMILES input strings was generated. For these heterocycle-substituted MAAs, only (hydroxy)cyclohexenone- and (hydroxy)palythine-type core structures were considered. Although such ring-containing substituents are somewhat atypical, they have been previously reported, for example in mycosporin-2 from the ascomycete *Gnomonia leptostyla* [51]. Furthermore, a linker moiety and a comprehensive set of building blocks were incorporated, including proteinogenic and non-proteinogenic amino acids, amine-containing osmolytes, as well as their decarboxylated derivatives. In contrast to the initial database, native substituents were also included in this iteration. The resulting combinatorial library, generated as a list of SMILES strings, was subsequently processed using the OPENBABEL chemical file format converter and imported into SIRIUS. While the first version of the database comprised 4,594 compounds with 1,233 unique molecular formulas, the updated CMAA-DB2.0 expands this space by approximately an order of magnitude. It now contains 44,880 compounds with 6,187 distinct molecular formulas, representing, to the best of our knowledge, the most comprehensive approximation of the theoretically accessible MAA chemical space.

**Figure 2.**
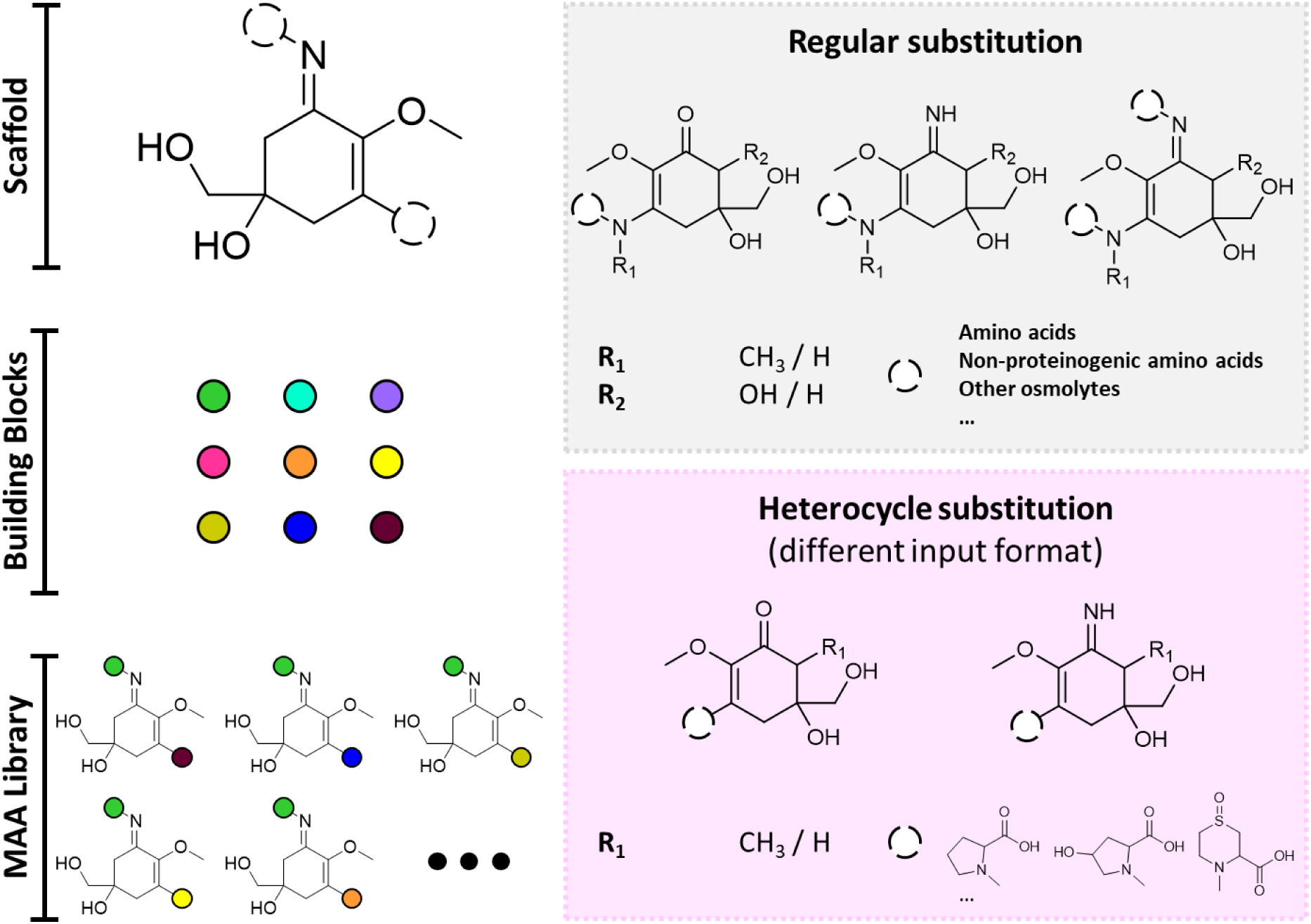
The theoretical concept for generating the combinatorial MAA database based on combining defined MAA scaffolds with a set of modular building blocks to enumerate all possible structures. In addition, two complementary strategies for subset generation are illustrated: one based on non-heterocyclic substituents, and another incorporating ring-containing substituents (e.g., (4-hydroxy)proline, chondrine, etc.).

To evaluate the applicability of the updated database (CMAA-DB2.0), the widely distributed and readily accessible red macroalga *Vertebrata lanosa* (Rhodomelaceae) was selected for our investigation. This seaweed is found along the coastal regions of Europe, North America, the Red Sea, the West Indies, and the Indo-Pacific [52]. It grows as an obligate epiphyte on the brown macroalga *Ascophyllum nodosum*. In non-scientific literature *V. lanosa* is often referred to as “ocean truffle“, and over generations it has been used as food and spice for locals living in coastal regions [53]. Lalegerie et al. previously identified several abundant MAAs, including shinorine, palythine, and asterina-330, alongside an unknown compound termed “MAA-1”, which was uniquely detected in this species among 40 investigated red macroalgae [54]. We therefore aimed to elucidate its identity. The crude methanolic extract of *V. lanosa* was analyzed using reversed-phase UHPLC-VWD-HRMS^2^, following established protocols from our previous studies [6, 20, 27, 55]. Application of our untargeted metabolomics workflow for MAA analysis, including data processing with mzmine, feature-based molecular networking on the GNPS2 web-platform, and metabolite annotation using SIRIUS, revealed an UV-active compound at a retention time of 1.16 min, hereafter referred to as compound **a**. In addition, several known and abundant MAAs were detected, including shinorine, palythine, asterina-330, porphyra-334, aplysiapalythine A, mycosporine-methylamine-threonine, usujirene, and palythene. Their absorption characteristics, exact masses of the corresponding proton adduct, and retention times were consistent with literature data and our previous analyses (Figure 3).

**Figure 3.**
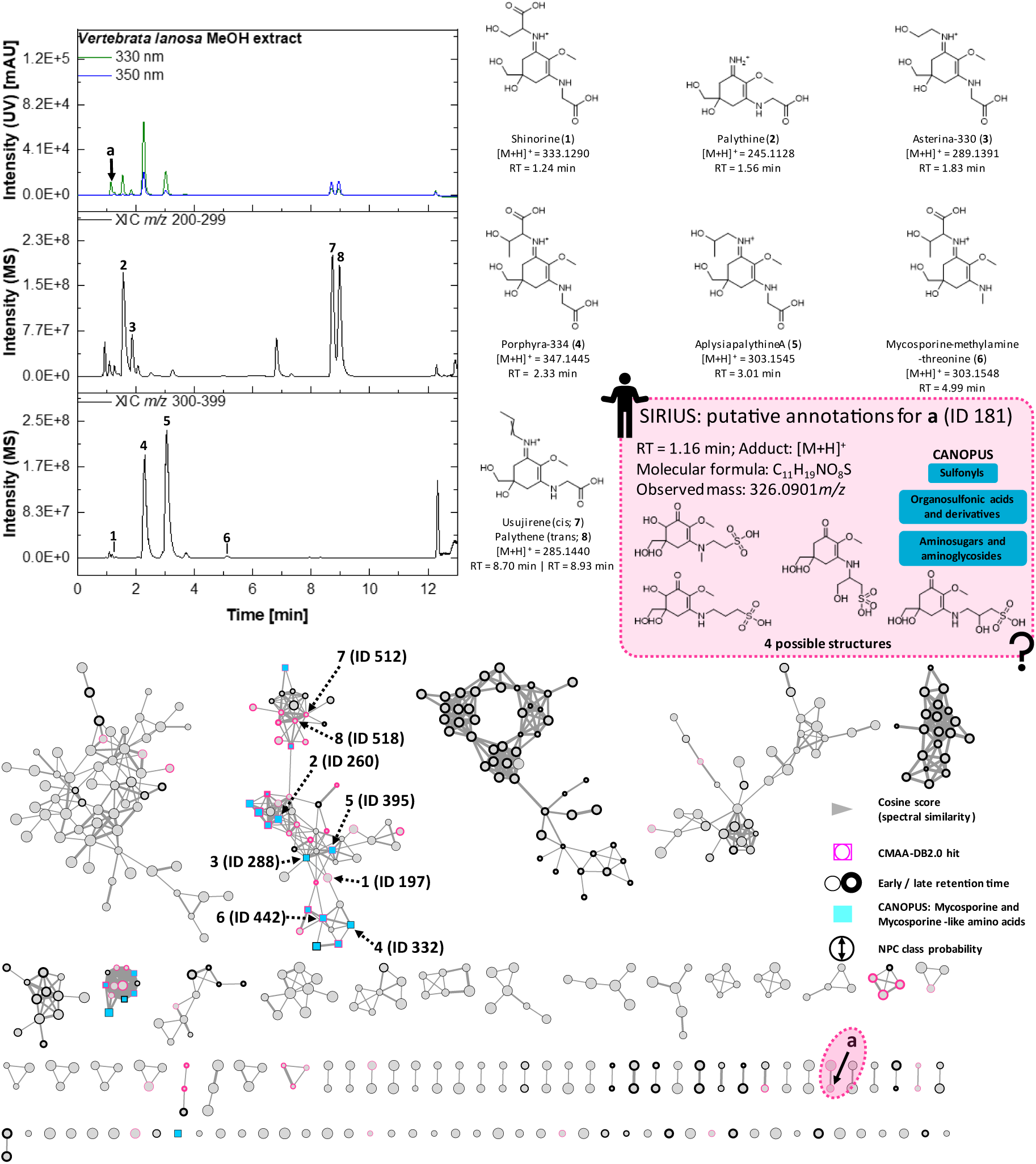
Chromatograms of a methanolic *Vertebrata lanosa* extract (UV detection at 330 and 350 nm; XICs *m/z* 200-299 and 300-399). Molecular structures of the [M+H]⁺ ions corresponding to putatively annotated MAAs (compounds 1-8) are shown together with the observed precursor ion masses and retention times. A feature-based molecular network generated from UHPLC-HRMS^2^ data of the *V. lanosa* extract and fractions S3-S7 is presented; some single nodes are not shown for clarity. The unknown compound **a** is highlighted by an arrow and a pink dashed circle; annotations and chemotaxonomic classification (SIRIUS) are provided. The following visual attributes were used: cosine score is represented by edge thickness; CMAA-DB2.0 hits by a pink node outline; retention time by node outline thickness; CANOPUS classification as mycosporines and mycosporine-like amino acids by cyan rectangles; and NPC class probability by node size.

In the feature-based molecular network generated via the GNPS2 workflow using cosine similarity scoring, all common MAAs were grouped within a single large cluster (Figure 3). For several features in this cluster, CANOPUS-based chemotaxonomic classifications as “Mycosporines and Mycosporine-like amino acids” at the NPC class level were obtained, along with corresponding hits from the CMAA-DB2.0. The unknown compound **a** (ID 181) was not associated with this cluster and instead appeared as a single node. This observation persisted when applying the DreaMS algorithm [56], an AI-based similarity scoring approach, as the feature remained unconnected to other putative MAAs (Figure S11B).

For the feature corresponding to compound **a**, SIRIUS predicted a molecular formula of C_11_H_19_NO_8_S and assigned the chemotaxonomic classes “Sulfonyls”, “Organosulfonic acids and derivatives”, and “Aminosugars and aminoglycosides”. Evidences supporting its classification as an MAA were its high polarity, characteristic UV absorption, and, most notably, the four candidate structures proposed by CMAA-DB2.0 (Figure 3). These comprised two MAAs featuring gadusol cores: one bearing *N*-methylation and a sulfonylethyl substituent, and the other a sulfonylpropyl substituent at the vinylogous amide functionality. In addition, a structure analogous to mycosporine-serinol, in which a hydroxyl group is replaced by a sulfonic acid moiety, was suggested, along with a cyclohexenone-type MAA carrying a 2-hydroxysulfonylpropyl substituent. All candidates represent previously undescribed natural products. However, the gadusol-core structures were considered less likely, as such compounds have thus far only been reported in the genera *Klebsormidium* [26] and *Apatococcus* [23]. In contrast, a mycosporine-serinol analogue and its constitutional isomer appeared more plausible. In the former case, the substituent corresponds to cysteinolic acid, a known osmolyte present in both bacteria and algae, including species of the genus *Vertebrata* [52]. Supporting this conclusion, a feature (ID 46) corresponding to the quasimolecular ion of cysteinolic acid was identified in the network (Figure S11A). For this feature, SIRIUS predicted the molecular formula C_3_H_9_NO_4_S + H^+^ (*m/z* 156.0323, Δ -1.28 ppm) and ranked the corresponding structure among the most probable candidates. To obtain definitive structural confirmation, targeted isolation of compound **a** was pursued using a combination of chromatographic techniques, including liquid-liquid extraction, reversed-phase MPLC, semi-preparative reversed-phase HPLC, and final purification by size-exclusion chromatography on Sephadex LH-20 material. Within the course of this isolation process, the known MAAs palythine, porphyra-334, asterina-330, and aplysiapalythine A were also obtained (Figure S1).

Compound **a** was obtained as a water-soluble, pale yellowish-white powder with high hygroscopicity. SIRIUS predicted for the corresponding feature (ID 181, [M+H]^+^ *m/z* 326.0901) the molecular formula C_11_H_19_NO_8_S (neutral molecule) with a median mass error of -0.43 ppm. The 2D-NMR data (Table 1, Figure 4) indicated a cyclohexenone-type scaffold bearing a methoxy group (δ_C_ 58.0 ppm, δ_H_ 3.42 s) at carbon C-2 and two methylene groups at C-4 (δ_C_ 33.1 ppm, δ_H_ 2.87 d, 2.33 d) and C-6 (δ_C_ 45.7 ppm, δ_H_ 2.29 d, 2.08 dd). Furthermore, one hydroxymethylene group (δ_C_ 67.7 ppm, δ_H_ 3.29 dd, 3.13 dd) was present, exhibiting HMBC correlations to C-4, C-5, and C-6. The spectrum was consistent with a vinylogous amide system extending from C-1 (δ_C_ 184.2 ppm, C=O) through C-2 (δ_C_ 129.7 ppm, C-OCH_3_) to C-3 (δ_C_ 152.4 ppm). In addition, two hydroxyl protons were observed at C-5 (δ_H_ 4.50 s) and C-7 (δ_H_ 4.88 m). The amide-associated proton (δ_H_ 6.49 d) showed HMBC correlations to C-2, C-4, and C-1’. The substituent was identified as cysteinolic acid, based on the presence of a methine group (δ_C_ 51.8 ppm, δ_H_ 3.77 m) as well as two methylene groups. Considerably different shifts of these methylene carbons (C-2’: δ_C_ 52.9 ppm; C-3’: 63.6 ppm) pointed towards varying substitutions. One hydroxyl group (δ_H_ 4.83 t) was found to be linked to C-3’, due to diagnostic HMBC cross-peaks to C-1’ and C-3’. Finally, a singlet at 8.50 ppm indicated a sulfonic acid substitution at C-2’. Tandem MS experiments supported this structural assignment, revealing a fragmentation pattern characteristic for MAAs, including two sequential losses of water, loss of the methoxy group, and cleavage of the cysteinolic acid substituent (Figure 4).

**Figure 4.**
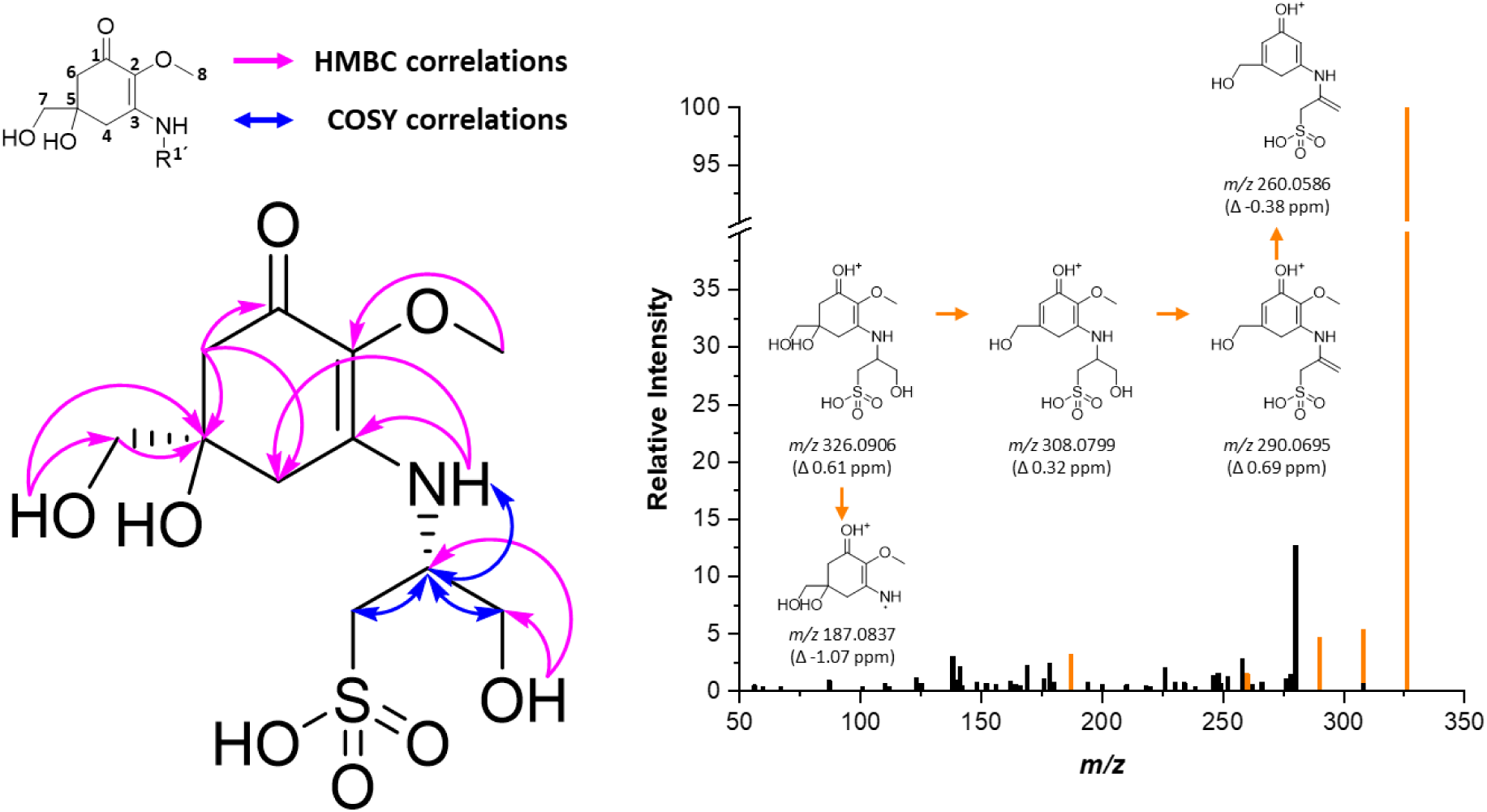
Molecular structure of mycosporine-cysteinolic acid (5R-1′R diastereomer) with key ^1^H-^13^C HMBC correlations indicated in pink and COSY correlations shown as blue arrows. The MS^2^ spectrum of mycosporine-cysteinolic acid ([M+H]^+^ ion, *m/z* 326.0906) is presented alongside. Peaks highlighted in orange represent characteristic MAA-specific fragment ions.

**Table 1.**
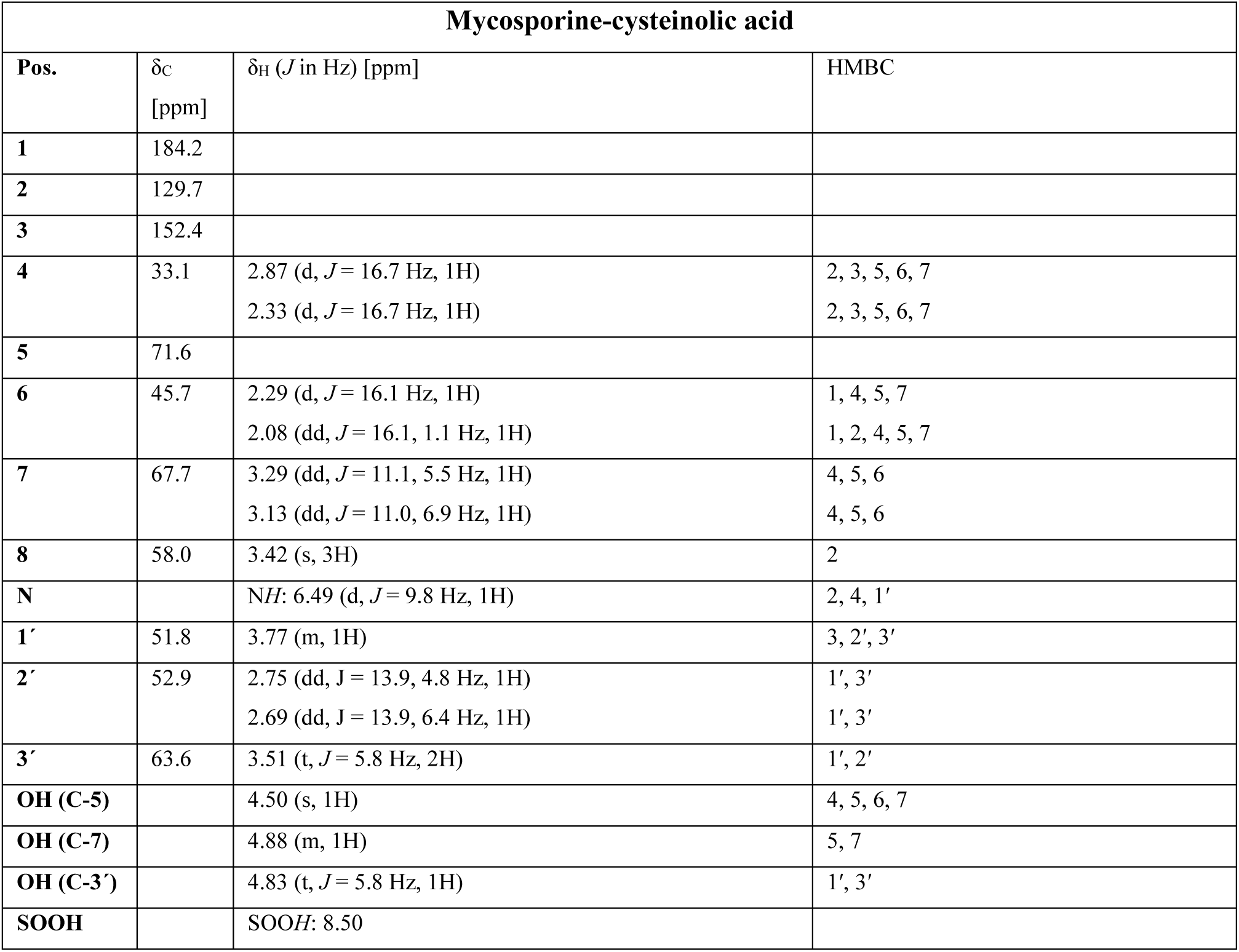
^1^H (600 MHz) and ^13^C (151 MHz) NMR data of mycosporine-cysteinolic acid. All spectra were recorded in DMSO-*d*_6_. For consistency with literature data, a numbering scheme was applied in which the *N*-substituent (side chain) was assigned the lowest possible position.

| Mycosporine-cysteinolic acid |  |  |  |
| --- | --- | --- | --- |
| Pos. | $\delta_C$<br>[ppm] | $\delta_H$ (*J* in Hz) [ppm] | HMBC |
| 1 | 184.2 |  |  |
| 2 | 129.7 |  |  |
| 3 | 152.4 |  |  |
| 4 | 33.1 | 2.87 (d, *J* = 16.7 Hz, 1H)<br>2.33 (d, *J* = 16.7 Hz, 1H) | 2, 3, 5, 6, 7<br>2, 3, 5, 6, 7 |
| 5 | 71.6 |  |  |
| 6 | 45.7 | 2.29 (d, *J* = 16.1 Hz, 1H)<br>2.08 (dd, *J* = 16.1, 1.1 Hz, 1H) | 1, 4, 5, 7<br>1, 2, 4, 5, 7 |
| 7 | 67.7 | 3.29 (dd, *J* = 11.1, 5.5 Hz, 1H)<br>3.13 (dd, *J* = 11.0, 6.9 Hz, 1H) | 4, 5, 6<br>4, 5, 6 |
| 8 | 58.0 | 3.42 (s, 3H) | 2 |
| N |  | NH: 6.49 (d, *J* = 9.8 Hz, 1H) | 2, 4, 1' |
| 1' | 51.8 | 3.77 (m, 1H) | 3, 2', 3' |
| 2' | 52.9 | 2.75 (dd, *J* = 13.9, 4.8 Hz, 1H)<br>2.69 (dd, *J* = 13.9, 6.4 Hz, 1H) | 1', 3'<br>1', 3' |
| 3' | 63.6 | 3.51 (t, *J* = 5.8 Hz, 2H) | 1', 2' |
| OH (C-5) |  | 4.50 (s, 1H) | 4, 5, 6, 7 |
| OH (C-7) |  | 4.88 (m, 1H) | 5, 7 |
| OH (C-3') |  | 4.83 (t, *J* = 5.8 Hz, 1H) | 1', 3' |
| SOOH |  | SOOH: 8.50 |  |

These findings, together with the SIRIUS-based annotation using the combinatorial MAA database, identify the unknown compound **a** in *V. lanosa* as a novel MAA, for which we propose the name mycosporine-cysteinolic acid. Polarimetry and circular dichroism experiments indicate that the compound is dextrorotatory, with the ring stereocenter at C-5 likely in S configuration. Since the naturally occurring isomer of cysteinolic acid is the D-form, and its conjugation with bile salts has been reported to preserve this configuration [57], it is plausible that the novel MAA has an S-configuration at C-1′. However, to elucidate the absolute configuration, we have resorted to theoretical calculations employing density functional theory (DFT). ECD spectra are known to be rather sensitive even to minor geometrical changes, hence we have computed the spectra by Boltzmann averaging over an ensemble of roughly 200 energetically low-lying conformations (see Section 3.7 Computational details). As Figure S11 shows, both the 5R-1′R and 5S-1′R diastereomers exhibit a prominent negative Cotton effect (positive peak at shorter wavelength, followed by negative peak at longer wavelength) observed in the experimental spectrum and centered around the absorption peak at 277 nm, the other two diastereomers reveal a positive Cotton effect and are hence excluded from further consideration. The theoretical spectral features are much narrower than the experimentally measured spectrum mainly due to the lack of vibronic coupling effects in our calculation.

Unfortunately, ECD alone does not provide sufficient information to decide between these two diastereomers. Thus, we have also computed the NMR chemical shifts of the carbon nuclei using DFT and an implicit water solvation model (Table S1). Both stereoisomers show relatively large errors (more than 2 ppm) in chemical shifts of the side-chain carbon atoms, in particular, atoms 8, 2′, and 3′. However, the absolute configuration at the 1′ center is the same in both molecules, and this deviation is expected due to the flexibility of these side chains on the one hand, and the lack of explicit solvation (the explicit presence of solvent water molecules) on the other hand, in particular the absence of the explicit solvation shell of the anionic sulfonate moiety. What is more revealing is the error in chemical shifts around the chiral center at position 5. Here it is clear that the conformationally-averaged chemical shifts of the 5R-1′R geometry is much closer (RMSD = 1.37 ppm) to the experimental values than the 5S-1′R (2.54 ppm). Hence, the DFT calculations of the ECD and NMR spectra, when considered together, indicate that the absolute configuration is 5R-1′R.

Furthermore, the compound exhibits a molar extinction coefficient of 6.3 x 10^3^ M^-1^ cm^-1^ at 309 nm in water, consistent with the high UV-absorbing properties typical for MAAs. To assess stability, an aqueous solution of the novel MAA was subjected to repeated freeze-thaw cycles. The sample was stored at -20 °C for 10 days with daily thawing, and HPLC-DAD monitoring demonstrated that the compound was stable under these conditions.

In a subsequent step, we investigated whether the novel MAA could also be detected in algal species other than *V. lanosa*. Table 2 represents the quantitative MAA profiles of seven selected compounds: mycosporine-cysteinolic acid (a), shinorine (1), palythine (2), asterina-330 (3), porphyra-334 (4), aplysiapalythine A (5), and the mixture of usujirene and palythene (6) in diverse algae. For quantification, a previously published HPLC-UV method was employed [29]. In total, nearly 40 different algal extracts were analyzed, including a commercially available *Porphyra* sp. sample, two *Vertebrata lanosa* samples (I and II), and previously unstudied algal samples from Peru. Interestingly, the results showed that MAA profiles varied considerably even among both *V. lanosa* samples. These variations appear to be influenced by both geographic location (approximately 55.9 km distance between the sampling locations Portsall and Roscoff in France) and harvest time, with *V. lanosa* (I) collected in August 2024 in Portsall and *V. lanosa* (II) collected in June 2018 in Roscoff. The most pronounced differences were observed for the novel MAA mycosporine-cysteinolic acid, which was nearly 20-fold higher in *V. lanosa* (I) compared to *V. lanosa* (II). Shinorine (1) was detected exclusively in *V. lanosa* (II), whereas the cis-trans isomers usujirene and palythene (6) were only present in *V. lanosa* (I). Considerable differences were also observed for asterina-330 (3), which was quantified at 1.14 mg/g dry weight in *V. lanosa* (I) but only 0.06 mg/g dry weight in *V. lanosa* (II). Overall, mycosporine-cysteinolic acid was only found in extracts of *V. lanosa*, which is in agreement with the findings of Lalegerie and colleagues [54].

**Table 2.** Quantitative MAA profiles in different algae as determined by HPLC-UV analysis. Mean values expressed as mg per g of DW biomaterial with corresponding absolute standard deviation. Compound assignment: mycosporine-cysteinolic acid (**a**), shinorine (**1**), palythine (**2**), asterina-330 (**3**), porphyra-334 (**4**), aplysiapalythine A (**5**), and usujirene & palythene (**6**). Det. = Detected but respective levels below LOQ.

|  | **a** | **1** | **2** | **3** | **4** | **5** | **6** |
| --- | --- | --- | --- | --- | --- | --- | --- |
| *Porphyra* sp. | - | 2.33 ± 0.30 | 0.30 ± 0.06 | 0.13 ± 0.04 | 18.10 ± 2.17 | 0.21 ± 0.13 | 0.55 ± 0.06 |
| *Vertebrata lanosa* (I) | 6.52 ± 0.50 | - | 1.87 ± 0.11 | 1.14 ± 0.04 | 2.48 ± 0.28 | 0.98 ± 0.18 | 0.06 ± 0.01 |
| *Vertebrata lanosa* (II) | 0.37 ± 0.02 | Det. | 1.26 ± 0.17 | 0.06 ± 0.01 | 1.02 ± 0.09 | 0.58 ± 0.02 | - |
| *Cryptopleura*<br>*cryptoneuron* | - | Det. | Det. | 0.24 ± 0.07 | - | - | - |
| *Griffithsia pacifica* | - | Det. | - | - | Det. | - | - |
| *Tiffaniella snyderae* | - | - | - | - | - | - | - |
| *Corallina chilensis* | - | 1.70 ± 0.05 | 0.82 ± 0.09 | 0.06 ± 0.02 | Det. | - | - |
| *Asterfilopsis centralis* | - | 8.96 ± 0.16 | 0.08 ± 0.03 | 0.03 ± 0.02 | Det. | - | - |
| *Sarcodiotheca*<br>*gaudichaudii* | - | 9.33 ± 0.51 | 0.08 ± 0.02 | 0.04 ± 0.01 | Det. | 0.14 ± 0.08 | - |
| *Iridaea tuberculosa* | - | 2.18 ± 0.18 | 2.79 ± 0.24 | 0.18 ± 0.04 | Det. | 0.17 ± 0.16 | - |
| *Chondrus*<br>*canaliculatus* | - | 3.23 ± 0.07 | 0.61 ± 0.03 | 0.07 ± 0.00 | - | - | - |
| *Chondracanthus*<br>*chamissoi* | - | 3.20 ± 0.02 | 1.47 ± 0.02 | 0.18 ± 0.02 | - | - | - |
| *Nitophyllum*<br>*peruvianum* | - | Det. | Det. | - | - | - | - |
| *Gracilariopsis*<br>*lemaniformis* | - | 1.75 ± 0.09 | 0.73 ± 0.06 | 0.48 ± 0.01 | 0.83 ± 0.03 | - | 0.11 ± 0.02 |
| *Prionitis decipiens* | - | 9.77 ± 0.07 | 1.43 ± 0.03 | 0.07 ± 0.00 | - | - | - |
| *Rhodomenia*<br>*flabellifolia* | - | - | - | - | - | - | - |

## 3 Materials and methods

### 3.1 Chemicals and solvents

Silica gel 40/63 (Merck, Darmstadt, Germany; product code 08965), Celite®545 (Merck; product code 102693), and Sephadex-LH20 (Merck; product code 17-0090-01) were used as column material. Thin-layer chromatography (TLC) was performed on aluminum-coated silica gel 60 plates (20 × 20 cm, 0.2 mm layer thickness, F254; Merck, product code 1.05554). HPLC-MS-grade solvents and chemicals included acetonitrile (Fisher Scientific, Loughborough, UK), methanol, formic acid (99–100%) (Merck), and ammonium formate (Sigma-Aldrich, St. Louis, MO, USA). Water was purified using an ARIUM 611 UV system (Sartorius, Göttingen, Germany). Extraction and TLC solvents (all analytical grade) comprised petroleum ether, dichloromethane, ethyl acetate, acetone, chloroform, acetic acid, and formic acid (all from VWR, Vienna, Austria), as well as methanol and 1-butanol (AnalaR NORMAPUR, VWR). The TLC spray reagent was prepared by dissolving 30 mg ninhydrin (SERVA Electrophoresis GmbH, Heidelberg, Germany) in 10 mL 1-butanol, assisted by ultrasonication, followed by addition of 0.3 mL acetic acid (98%). For NMR analysis, DMSO-*d*_6_ (Merck; product code 1003645030) was used as solvent.

### 3.2 General instrumentation

A Bosch MKM 6003 grinder (Stuttgart, Germany) was used to homogenize biomaterial. Ultrasound-assisted extraction was conducted using a Sonorex TK 52 ultrasonic bath (Bandelin, Berlin, Germany), and evaporation of solvents with a LABOROTA 4000 rotary evaporator (Heidolph, Schwabach, Germany). Freeze-drying was performed in a Benchtop Pro (SP Scientific, Warminster, PA, USA) device, for centrifugation, a Labofuge 400 (Heraeus, Hanau, Germany) or a Centrifuge 5804R (Eppendorf, Hamburg, Germany) were used. A Vortex-Genie 2 (Scientific Industries, Bohemia, NY, USA) was used for mixing solutions. A Reveleris® X2 system (BÜCHI, Flawil, Switzerland), equipped with a binary pump, an autosampler, an automated fraction collector, and an evaporative light-scattering detector (ELSD) as well as UV detector was employed for reversed-phase medium-pressure chromatography (RP-MPLC) separation. Semi-preparative HPLC was carried out on an Agilent Technologies 1260 Infinity II system (Santa Clara, CA, USA) consisting of a quaternary pump (G7157A), a column oven (G7116A), a variable wavelength detector (G7114A), and a fraction collector (G1364F9). Solutions were transferred with pipettes and tips from Eppendorf. HPLC analyses were carried out on a Merck-Hitachi Elite LaChrom system (Tokyo, Japan) equipped with a quaternary pump, vacuum degasser, autosampler, column thermostat, and diode-array detector. NMR experiments were conducted on spectrometers from Bruker (Karlsruhe, Germany): 400 MHz Avance NEO (^1^H: 400 MHz; ^13^C: 101 MHz) and 600 MHz Avance NEO (^1^H: 600 MHz; ^13^C: 151).

### 3.3 Origin of biomaterial, sample preparation, and isolation of MAAs from *Vertebrata lanosa*

Commercially available *Porphyra* sp. biomaterial was bought in a local supermarket in Innsbruck 2026. *Vertebrata lanosa* (I) was collected in Portsall, near Ploudalmézeau, France and morphologically authenticated by Solène Connan (UBO, Université de Bretagne Occidentale). *Vertebrata lanosa* (II) was collected in Roscoff, France and morphologically identified by Prof. Ulf Karsten (University of Rostock, Germany). Further red algae were collected in the Paracas District, Pisco Province, Ica Department, Peru, by the research team of the Laboratorio de Investigación en Cultivos Marinos (LICMA) and were morphologically identified by specialists at the Natural History Museum of the Universidad Ricardo Palma. Voucher specimens of all samples are deposited at the Department of Pharmacognosy, University of Innsbruck, Austria. For additional information see Table S1.

Approximately 2 kg of dried *Vertebrata lanosa* biomass was sequentially extracted with solvents of increasing polarity (petroleum ether, dichloromethane, ethyl acetate, methanol, methanol/water 5:1 v/v, and water), yielding 2.66 g, 7.35 g, 1.43 g, 167.71 g, 132.87 g, and 51.90 g of dried extracts, respectively. The methanol (FM) and methanol/water (FH) extracts, enriched in mycosporine-like amino acids (MAAs), were further purified via sequential liquid-liquid extraction with ethyl acetate and *n*-butanol to remove remaining apolar compounds and polysaccharides. The combined FM and FH extracts (FMH) were fractionated by silica gel column chromatography using an ethyl acetate-methanol gradient (S1-S7). Fractions were evaluated by TLC (*n*-butanol:acetone:acetic acid:water, 10:4:2:0.5 v/v/v/v) and HPLC-UV [29] analysis. Obtained collection fractions were subsequently purified using a combination of RP-MPLC (C18 column, water-methanol-acetonitrile gradient), semi-preparative RP-HPLC, and Sephadex LH-20 chromatography (MeOH) to isolate individual MAAs. Fractions were monitored by HPLC-UV [29], TLC, and UPLC-MS [58], pooled according to their composition, and yields recorded as detailed in the isolation tree (Figure S1). Experimental details regarding extraction and isolation are depicted in the supplementary information (Sections S1.1.-S1.5.).

### 3.4 Untargeted metabolomics analyses by UHPLC-VWD-ES-HRMS/MS and feature-based molecular networking

UHPLC-VWD-HRMS^2^ analyses were performed on a Vanquish system coupled to a variable wavelength detector and a Thermo Scientific Exploris 120 Orbitrap HRMS unit (Thermo Scientific, Waltham, MA, USA), as described previously [23]. Chromatographic separation was achieved on a Luna Omega C18 100 Å column (100 mm × 2.1 mm; particle size: 1.6 μm; Phenomenex, Torrance, CA, USA), equipped with a SecurityGuard ULTRA guard C18 pre-column containing the same stationary phase. The mobile phase consisted of water containing 0.25% formic acid and 20 mM ammonium formate (A) and acetonitrile (B), applied in a gradient elution over a total runtime of 20 min. Raw MS data were converted using MSConvert [59] and subsequently processed in mzmine (4.8.30) (mzio GmbH, Bremen, Germany) [60] using the processing wizard for batch generation and applying the following parameters. Global settings: The preset UHPLC-Orbitrap-DDA was selected. Smoothing and stable ionization across samples were enabled. The retention time (RT) range was cropped to 0.3-13 min. The maximum number of peaks per chromatogram and the minimum number of consecutive scans were set to 15 and 5, respectively. The approximate feature FWHM (full-width-at-half-maximum) was set to 0.2 min, while RT tolerances were defined as 0.05 min (intra-sample) and 0.25 min (sample-to-sample). Orbitrap settings: The ionization mode was set to positive ESI mode. Absolute intensity noise thresholds were defined as 7 x 10^5^ for MS^1^ and 0 for MS^2^. The minimum feature height was set to 7 x 10^5^. The scan-to-scan *m/z* tolerance was 0.0012 (4 ppm), the intra-sample *m/z* tolerance 0.0015 (5 ppm), and the sample-to-sample *m/z* tolerance 0.0021 (7 ppm). Filters: The original feature list was removed, and only features containing ^13^C isotopes were retained. Annotation: A local compound database search was performed using an in-house library of mycosporine-like amino acids (*m/z* tolerance: 0.0021 or 7 ppm; https://zenodo.org/records/21775642). Additionally, the lipid annotation module was enabled. DDA: The export functions for molecular networking and SIRIUS were activated. After batch file generation, an additional “Feature list rows filter” module was inserted between the “Duplicate peak filter” and “Correlation grouping” steps to reset feature IDs and retain only rows containing MS^2^ spectra. All molecular networks were visualized using Cytoscape software [61].

The DreaMS [56] network was created using the Spectral / Molecular Networking function within mzmine with the following settings: merge & select fragment scans, merged (simple); presets, single scan: merged across energies; merging *m/z* tolerance: 0.0021 *m/z* or 7.0 ppm; min similarity: 60%; k-nearest neighbors, num. neighbors = 3, min neighbor similarity = 60%; batch size, 32 [56].

The feature-based molecular network (FBMN) [62, 63] was created on the GNPS2 webpage (https://gnps2.org/homepage) [64] with the mzmine-processed data: The precursor ion mass tolerance was set to 0.02 Da and the MS/MS fragment ion tolerance to 0.02 Da. Both, the window filter as well as the precursor window filter were allowed. A molecular network was then created where edges were filtered to have a cosine score above 0.6 and more than 5 matched peaks. Further, edges between two nodes were kept in the network if and only if each of the nodes appeared in each other’s respective top 10 most similar nodes. The maximum size of a molecular family was set to 100, and the lowest scoring edges were removed from molecular families until the molecular family size was below this threshold. The spectra in the network were then searched against GNPS spectral libraries, with the analog search function activated. The library spectra were filtered in the same manner as the input data. All matches kept between network spectra and library spectra were required to have a score above 0.6 and at least 4 matched peaks. Additional edges were provided as well. The FBMN job can be publicly accessed at: https://gnps2.org/status?task=976145b48c57402ea3346038327c9e90.

### 3.5 Generating the new combinatorial MAA database (CMAA-DB2.0)

The combinatorial MAA database was created using the free software SmiLib v2.0 (http://melolab.org/smilib/) [50]. For the first subset, which included heterocyclic or secondary amine building blocks such as proline or rhodoic acid, four scaffolds were employed: cyclohexenone, hydroxycyclohexenone, palythine, and hydroxypalythine. The second subset, featuring regular substitution patterns, was generated using twelve different scaffolds, including the *N*-methyl, *N*-methylhydroxy, and hydroxy derivatives of the cyclohexenone, palythine, and aminocyclohexenimine cores. For all inputs, the so-called empty linker [A][R1] was used. Building blocks included native and decarboxylated (non-)proteinogenic amino acids, as well as other organic osmolytes. Different representations of the building blocks were used depending on the subset. The fully enumerated libraries were saved as .txt files and subsequently converted to a different SMILES code representation using the OPENBABEL chemical file format converter (https://www.cheminfo.org/Chemistry/Cheminformatics/FormatConverter/index.html). The consolidated, converted list was then saved as a .tsv file and used to generate a SIRIUS-compatible combinatorial MAA database using the dedicated “Import Custom Databases” function in SIRIUS [65]. In total, the database comprised 44,880 compounds with 6,187 unique molecular formulas.

### 3.6 SIRIUS metabolite annotation

Metabolite annotation was performed with SIRIUS 6.3.3 [65–67]. The corresponding .mgf file was exported from mzmine and subsequently processed. The parameters were set as follows: instrument, Orbitrap; MS^2^ mass accuracy, 10 ppm; fix formula for detected lipid, yes; fallback adducts: [M+H]^+^, [M+Na]^+^, and [M+K]^+^; search DBs, CMAA-DB2.0 (i.e., the new combinatorial MAA database). The spectral matching module was enabled to compute the similarity between all experimental features with the selected database (identity search, 10 ppm precursor deviation; analogue search enabled). The SIRIUS molecular formula identification was used with the de novo + bottom up strategy (default settings). The prediction of fingerprints was carried out with CSI:FingerID and the prediction of chemical classes with CANOPUS.

### 3.7 Computational details

All calculations were performed in Orca version 6.1.1 [68].

#### 3.7.1 Conformational analysis

Since ECD spectra are known to be rather sensitive even to minor geometrical changes, we have generated an ensemble of conformations for both the 5S,1’S and 5S,1’R stereoisomers, which are referred to as the SS and SR stereoisomer, respectively.

For calculation, the global optimization algorithm GOAT [69] in combination with the semi-empirical potential GFN2-xTB [70] and the ALPB implicit solvation model for water [71] were used.

For the SR stereoisomer, in addition to the global energy minimum, 172 conformations were found within an energy window of 6 kcal/mol from the global minimum. 255 conformers were found in the same energy window relative to the SS global minimum. The conformers of the SS and RS stereoisomers, when needed, were obtained by inverting the coordinates of the RR and SR isomers, respectively.

Each of the generated conformations was re-optimized with density functional theory (DFT) using the range-separated hybrid functional wB97X [72] and the Ahlrich triple-Zeta Def2-TZVP basis set with the CPCM water model [73] and D4 dispersion correction [74]. The vibrational frequencies of the optimized geometries were also computed at the same level, and it was verified that all the obtained geometries are indeed energy minima. The obtained free energies were used to compute the statistical weight of each conformer according to the Boltzmann distribution.

#### 3.7.2 Computing the electronic circular dichroism (ECD) spectra

The ECD spectra of all conformers were computed using the same DFT setup and time-dependent density functional theory (TDDFT). Twelve roots were computed, and to improve the predicted intensity of the transitions, the Tamm-Dancoff approximation was omitted. The spectra were finally averaged using the Boltzmann weights. The spectra for the RR and RS stereoisomers were obtained via mirror reflection of the SS and SR spectra relative to the x-axis. As is usually necessary in TDDFT, the spectra were slightly shifted, by +15 nm, to match the experimental spectrum.

#### 3.7.3 Computing the NMR nuclear magnetic shieldings and chemical shifts

Nuclear magnetic shieldings of the carbon and hydrogen nuclei in the RR and SR conformers were computed with DFT using the TPSS meta-GGA functional [75] and the segmented contracted triple-Zeta basis set pcSseg-2 [76], also with CPCM solvation in water. The computed values for all the conformations were Boltzmann-averaged, and then shifted relative to the shieldings of tetramethylsilane (TMS) computed with the same setup, to finally obtain the conformationally-averaged chemical shifts. The reference magnetic shielding values from TMS were 184.83 ppm for carbon and 31.74 ppm for hydrogen.

### 3.8 MAA screening study of different algal extracts

Algal biomaterial was milled and sieved (180 µm). Subsequently, 10 mg of each sample was weighed into a 1.5 mL reaction tube. Extraction was performed three times using 1 mL of 25% MeOH as solvent, with 15 min of ultrasound-assisted maceration between each extraction step. After each cycle, the samples were centrifuged (15,000 rpm) for 5 min. The supernatants were combined, transferred into small glass vials, frozen at -80 °C, and subsequently subjected to lyophilization. The dry residue was re-dissolved in 400 µL of pure water and filtered through a cotton-plugged Pasteur pipette directly into HPLC vials prior to analysis. All samples were analyzed in triplicate. Quantification was performed with the method of Orfanoudaki et al. [29]. For mycosporine-cysteinolic acid, seven concentrations in the range of 7.81–500 µg/mL (solvent water) were prepared and analyzed. The calibration curve exhibited a linear relationship described by the regression equation y = 27.814x + 37.1, with a coefficient of determination (R²) greater than 0.999. The limit of detection (LOD) and limit of quantification (LOQ) were determined to be 1.25 µg/mL and 3.79 µg/mL, respectively.

## 4 Conclusions

Mycosporine-like amino acids remain an active area of research, with novel compounds still being discovered even in already well-studied organisms, as demonstrated here by the first report of mycosporine-cysteinolic acid in *Vertebrata lanosa*. The application of the new and significantly expanded database CMAA-DB2.0 enabled the annotation of this compound prior to any preparative isolation and greatly facilitated structure elucidation through NMR and MS experiments. Beyond its role in identifying novel compounds in algal extracts, the database also supports studies on MAA biosynthesis by allowing targeted searches for specific building blocks within organisms. Its modular design offers additional flexibility, enabling users to select which scaffolds to include. While the full database may not be required for certain red algae, which likely lack algasporine-type MAAs, it allows comprehensive screening of experimental MS^2^ spectra against more than 44,000 combinatorial MAA structures when broader coverage is desired. The alga *V. lanosa* proves to be a valuable model for applying this strategy in order to explore algal MAA diversity, as samples from geographically close locations exhibited substantial quantitative differences. This underscores the potential influence of collection site, harvest time, and extraction or isolation procedures on composition. Beyond facilitating the discovery of previously unrecognized MAAs, the expanded CMAA-DB2.0 provides a versatile resource for future metabolomics-guided investigations of MAA diversity across taxonomically and ecologically diverse organisms. As additional, experimentally validated MAA structures become available from time to time, the database can be continuously refined and expanded, thereby improving annotation confidence and enabling an increasingly comprehensive exploration of MAA biosynthesis and structural diversity.

In conclusion, integrating a substantially expanded combinatorial library with established preparative chromatographic workflows and advanced structure elucidation techniques resulted in an improved platform for MAA identification. This approach helped to annotate known and novel MAAs and provides a robust framework for accelerating future discoveries and advancing our understanding of MAA chemistry and biosynthesis.

## 5 Acknowledgements

The project under which the species *Chondracanthus chamissoi* and *Gracilariopsis lemaneiformis* were obtained was registered under code PRE-5-2023-00455, while the remaining Peruvian rhodophyte species were obtained under Project Code No. 018-2025-PRO99. Ethical approval for the research procedures was granted by the Ethics Committee of Universidad Científica del Sur (*Constancia de Comité de Ética* No. 126-CIEI-AB-CIENTÍFICA-2023). Paul Baltazar is acknowledged for facilitating access to biological material in coordination with the Laboratorio de Investigación en Cultivos Marinos (LICMA) and CONTRAPALMAR. Access to genetic resources and their derivatives was authorized by the Ministry of Production of Peru (PRODUCE) under *Resolución Directoral* No. 00109-2025-PRODUCE/DGAAMPA. Official taxonomic identification of all biological samples of Peruvian origin was conducted at the Natural History Museum of Universidad Ricardo Palma under *Constancia de Determinación Taxonómica* No. 001-2026-MURP. S.C. would like to thank Sylvain Petek, Alain Guenneguez, and Marie-Aude Poullaouec (LEMAR, Univ. Brest) for their help in collecting and grinding the 20 kg of *Vertebrata lanosa* from Portsall. The authors gratefully acknowledge the computing time provided to them at the Paderborn Center for Parallel Computing PC2.

## 7 Author information

Armin Oberosler and Fabian Jürgen Hammerle have contributed equally to this work and share first authorship.

### 7.1 Contributions

A.O.: Investigation, Formal analysis, Data curation, Writing – Original Draft, Writing – Review & Editing. F.J.H.: Conceptualization, Investigation, Formal analysis, Data curation, Writing – Original Draft, Writing – Review & Editing, Visualization, Supervision. S.L.: Investigation, Formal analysis. H.E.: Investigation, Formal analysis, Writing – Original Draft, Writing – Review & Editing. S.C.: Conceptualization, Resources, Writing – Review & Editing. F.P.: Resources, Writing – Review & Editing. B.B.: Resources, Writing – Review & Editing. U.K.: Resources, Writing – Review & Editing, Funding acquisition. M.G.: Resources, Writing – Review & Editing, Supervision, Project administration, Funding acquisition.

### 7.2 Corresponding author

Correspondence to Fabian Jürgen Hammerle.

## 8 Ethics declarations

### 8.1 Consent for publication

All authors have read this manuscript and would like it to be considered for publication.

