## Supplementary information for "Discovery of a novel UV-absorbing mycosporine-like amino acid in *Vertebrata lanosa* using an expanded combinatorial structure database incorporating non-proteinogenic amino acids and organic solutes"

Armin Oberosler[1]*, Fabian Jürgen Hammerle[1]*°, Sebastian Lanner[1], Hossam Elgabarty[2], Solène Connan[3], Fernanda Pita[4,5], Barış Ballık[5], Ulf Karsten[6,7], Markus Ganzera[1]

* Authors contributed equally

° Correspondence

***Affiliations***

[1] Institute of Pharmacy, Department of Pharmacognosy, Center for Chemistry and Biomedicine, University of Innsbruck, 6020 Innsbruck, Austria

[2] Department of Chemistry, Paderborn University, 33098 Paderborn, Germany

[3] Univ Brest, CNRS, IRD, Ifremer, LEMAR, 29290 Plouzane, France

[4] Grupo de Investigación en Acuicultura Sostenible, Facultad de Ciencias Ambientales, Universidad Científica del Sur, Lima 15842, Peru

[5] Department of Cell Biology of Phototrophic Marine Organisms, Institute of Biosciences, University of Rostock, 18059 Rostock, Germany

[6] Department of Applied Ecology and Phycology, Institute of Biosciences, University of Rostock, 18059 Rostock, Germany

[7] Interdisciplinary Faculty, Department of Maritime Systems, University of Rostock, 18059 Rostock, Germany

### 1. Extraction and isolation of mycosporine-like amino acids (MAAs) from *Vertebrata lanosa*

S1.1. Extraction

2015.6 g of dried *Vertebrata lanosa* (I) biomass was sequentially extracted using solvents of increasing polarity. One liter of each solvent was used for extraction unless stated otherwise. Algal material and solvents were shaken for 3 min and subsequently subjected to ultrasound-assisted extraction for 15 min at room temperature. After maceration, the solutions were filtered using a vacuum filtration flask, followed by a second filtration step with a paper filter (particle retention capacity: 5–13 µm). The filtrates were evaporated to dryness under reduced pressure using a rotary evaporator with a water bath temperature of 40 °C. Each extraction procedure was performed in quadruplicate unless stated otherwise. The final extracts were transferred into pre-weighed, sealable containers and dried under a stream of compressed air. Extraction with dichloromethane was performed six times instead of four, and methanol extraction was also conducted six times. Extraction with methanol and water (8:2 v/v) was performed three times. For the extraction using methanol and water (1:1.5 v/v), 1.0 L of methanol and 1.5 L of water were used. The resulting extracts were designated as FP (petroleum ether), FDCM (dichloromethane), FEAc (ethyl acetate), FM (methanol), FH (methanol/water 5:1 v/v), and Fx (water). The yields of the dried extracts were 2.66 g (0.13 %) for FP, 7.35 g (0.36 %) for FDCM,

1.43 g (0.07 %) for FEAc, 167.71 g (8.32 %) for FM, 132.87 g (6.59 %) for FH, and 51.90 g (2.57 %) for Fx. The weight of the residual biomaterial was 1,522.30 g (75.53 %). The two MAA-containing extracts, FM and FH, were further purified. The extracts were dissolved in HPLC-grade water and (separately) subjected to liquid-liquid extraction with ethyl acetate and *n*-butanol. Liquid-liquid extraction of FM yielded 136.30 g (81.27%) of the aqueous fraction (FM-water), 24.43 g (14.57%) of the ethyl acetate fraction (FM-EAc), and 1.44 g (0.86%) of the *n*-butanol fraction (FM-but). For FH, the extraction yielded 106.08 g (80.38%) of the aqueous fraction (FH-water), 8.77 g (6.60%) of the butanol fraction (FH-but), and 6.33 g (4.76%) of the ethyl acetate fraction (FH-EAc). FM-water and FH-water were then combined into a single fraction, designated FMH, which was subsequently used for further separation and purification of MAAs.

S1.2. Silica gel column chromatography

Silica gel column chromatography of the dried fraction FMH was performed using an ethyl acetate to methanol gradient. The column (diameter = 13.5 cm, length = 28 cm) was packed with 1.5 kg of silica gel (particle size 400-630 µm) (Merck KGaA, Darmstadt, Germany). Using a mortar and pestle, 200.0 g of dried FMH was finely ground and passed through a sieve with a pore size of 355 µm. The sieved fraction was then triturated with silica gel (200.0 g, particle size 400-630 µm). Elution was carried out with 2 L each of 100:0, 80:20, 60:40, and 40:60 (v/v) ethyl acetate-methanol, followed by 3 L of 20:80 and 5 L of 0:100 (v/v) methanol, resulting in a total elution volume of 16 L. The flow rate was kept between 3 mL/min and 5 mL/min and eluates were collected in 250 mL Erlenmeyer flasks manually. For the last three fractions, N1-N3, approximately 2 L of eluate was collected for each fraction. Vials 1–20 (S1, 7085.88 mg) were pooled, as no MAAs were observed within these fractions. Vials 21 and 22 were combined into S2 (7383.64 mg), vials 23–26 into S3 (11870.25 mg), vials 27–29 into S4 (13973.1 mg), and vials 30–32 into S5 (10821.52 mg). The vials 33–38 and fraction N1 were pooled together to form S6 (14857.1 mg). Finally, S7 (41321.98 mg) was formed from factions N2 and N3. Pooling was decided upon both TLC (mobile phase: *n*-butanol:acetone:acetic acid:water, 10:3:2.5:1 v/v/v/v) and HPLC-UV analysis of each obtained fraction.

S1.3. Reveleris® X2 RP-MPLC

8.03 g of fraction S4 and 15.00 g of fraction S7 were used and triturated with Celite545® (Merck KGaA, Darmstadt, Germany) in a 1:1 ratio. These mixtures were subjected to reversed-phase medium-pressure liquid chromatography (RP-MPLC) separation under following conditions: A Reveleris® X2 instrument from Grace/Büchi (BÜCHI Labortechnik AG, Flawil, Switzerland), which is equipped with an autosampler as well as both an ELS and a UV detector, was used. A $C_{18}$ cartridge from Büchi (BÜCHI Labortechnik AG, Flawil, Switzerland) was used as the stationary phase (pore size: 40 µm, weight: 40 g). For fraction S4, the flow rate was set at 5 mL/min and the runtime was 90 min. The detection wavelengths were set to 254 nm and 330 nm. The per-vial volume was set at 15 mL, and the non-peak volume was adjusted to 20 mL. For the separation of fraction S7, the flow rate was 10 mL/min. All other parameters were identical. The elution started with isocratic conditions of 100% water from 0 to 50 min. Between 50 and 70 min, the mobile phase composition was linearly changed to 100% methanol, which was then maintained for 10 min. At 80.1 min, the eluent was switched to 100% acetonitrile and held for 10 min. Fractionation of S4 resulted in three collection fractions: S4R1 (vials 2–5, 5600.63 mg), S4R2 (vials 6–11, 4024.01 mg), and S4R3 (vials 12–57, 1663.64 mg) (Figure S1). Fraction S4R2 contained the highest concentration of UV active compounds (potentially MAAs), therefore further purification was carried out with this fraction. Fractionation of S7 resulted in five collection fractions: S7R1 (vials 1–11, 7518.28 mg), S7R2 (vials 12–18, 1835.6 mg), S7R3 (vials 19–21, 93.57 mg), S7R4 (vials 22–24, 673.98 mg), and S7R5 (vials 35–end, 489.63 mg). Fraction S7R2 contained the highest concentration of UV active compounds (potentially MAAs), therefore further purification was carried out with this fraction.

S1.4. Semi-preparative HPLC

Fractions S4R2 (4024.01 mg) and S7R2 (603.85 mg) were dissolved in 4 and 2 mL HPLC-grade water respectively and (separately) subjected to semi-preparative RP-HPLC. Following conditions were employed: an Agilent (Agilent Technologies, Inc., Santa Clara, CA, USA) semi-preparative 1260 Infinity II HPLC system was used, including an autosampler, a quaternary pump, a variable-wavelength detector, and a fraction collector. The entire process was controlled using the Agilent Prep LC 3D ChemStation software. The detection wavelengths were set to 300 nm and 330 nm. A YMC (YMC Co., LTd., Kyoto, Japan) Actus Triart C18 110 Å (150 × 10 mm; 5 µm) was used as the stationary phase. The injection volume was set to 25 µL, the flow rate to 2.7 mL/min, and the temperature to 20 °C. The separation was performed using 0.1% formic acid in water (A) and methanol (B) as mobile phases. The gradient program was as follows: 0–20 min, 100% A; 20–30 min, linear gradient to 80% A; 30–35 min, linear gradient to 2% A; 35–40 min, 2% A; 40–55 min, re-equilibration with 100% A. The total run time was 55 min. Collected fractions were frozen at -80 °C and subjected to freeze drying. For

S4R2, fractions were collected based on retention time as follows: S4R2SP1 (14.9–16.1 min, 730.53 mg), S4R2SP2 (16.2–18.0 min, 98.9 mg), S4R2SP3 (19.5–22.6 min, 233.63 mg), S4R2SP4 (24.5–26.8 min, 15.72 mg, **asterina-330**), S4R2SP5 (37.4–38.2 min, 80.09 mg, **aplysiapalythine A**), S4R2SP6 (38.3–38.5 min, 13.57 mg), S4R2SP7 (38.6–39.6 min, 5.01 mg), S4R2SP8 (39.7–40.5 min, 3.26 mg), and S4R2SP9 (43.6–44.0 min, 20.47 mg).

Fraction S7R2 was separated on a Phenomenex® (Phenomenex Inc., Torrance, CA, USA) Synergi 4u POLAR-RP 80 Å (250.00 x 10.00 mm; 4 µm) column. Column oven temperature and injection volume were set to 5 °C and 75 µL, respectively. The chromatographic separation was performed using the following gradient program (solvent A was 0.1% formic acid in HPLC-grade water; solvent B was MeOH): 0–14 min, 100% A; 14–15 min, linear change to 1% A; 15–18.5 min, 1% A; 18.5–18.6 min, 100% A; 18.6–31 min, re-equilibration with 100% A. Collected fractions were frozen at -80 °C and subjected to freeze drying. For S7R2, fractions were collected based on retention time as follows: S7R2SP1 (7.22–7.79 min, 5.91 mg), S7R2SP2 (8.96–10.95 min, 196.02 mg, **porphyra-334**), Fraction S7R2SP3 (11.43–13.02 min, 6.55 mg).

S1.5. Sephadex LH-20 column chromatography of fraction S4R2SP1

A glass column (diameter = 25 mm, length = 750 mm) was packed with Sephadex LH-20 material (Merck KGaA, Darmstadt, Germany). The column was loaded with 167.84 mg of fraction S4R2SP1, which was dissolved in 1 mL of MeOH and filtered through a Pasteur pipette with a cotton plug. For fraction collection, an LKB (LKB-Produkter AB, Bromma, Sweden) Bromma 2211 100 Superrac was employed with racks of the type B. The flow rate was 2 mL/min and 4 mL per test tube were collected. After the collection process, each tube was analyzed by TLC, HPLC-UV, and UPLC-MS [1]. Pooling of fractions was done in accordance with the analytical results. Collected fractions were dried with a gentle stream of air, redissolved in HPLC-grade water, frozen at -80 °C, and subjected to freeze drying. Fractionation of S4R2SP1 resulted in following collection fractions: S4R2SP1SEC1 (vials 1–20, 0.21 mg), S4R2SP1SEC2 (vials 21–40, 115.28 mg), S4R2SP1SEC3 (vials 41–45, 3.4 mg, **mycosporine-cysteinolic acid**), and S4R2SP1SEC4 (vials 46–50, 0.78 mg).

#### 2. Physicochemical characterization of mycosporine-cysteinolic acid

S2.1. Polarimetry

A P-2000 polarimeter from Jasco (JASCO Corporation, Tokyo, Japan) with the serial number B184861232 was utilized. Flint glass was used as the Faraday cell. The light source was a sodium lamp operating at a wavelength of 589 nm. Ten measurement cycles were performed, from which the mean value and standard deviation were calculated. The temperature was set to 20 °C. A measurement tube from Jasco (Type J/39.35; length = 100 mm) was used. First, the solvent was measured as a blank. Then, the individual sample was filled into the analysis tube and measured. The blank value was subtracted from the measured sample value. The sample was measured at a concentration of 2.33 mg/mL in water. 1 mL of sample solution was used for measurement.

S2.2. Circular dichroism

A solution of mycosporine-cysteinolic acid was prepared at a concentration of 0.21 mg/mL in water. After preparing the solution, a water blank was measured first. A Jasco J-1500 CD spectrometer including a thermostatic unit (Jasco CTU-100, serial number B 026661871) was employed for the measurements. A cuvette made from high-performance quartz glass from Hellma Analytics (Hellma GmbH & Co. KG, Müllheim, Germany) (serial number 104-10-40) with a light path of 10 mm was used. The temperature was set to 25 °C and the CD-spectrum was recorded using SpecDis v.1.7 software (SpecDis, Berlin, Germany).

S2.3. Infrared spectroscopy

An infrared spectral measurement was performed on Alpha II FT-IR spectrometer of the manufacturer Bruker (Bruker Corporation, Billerica, MA, USA). Bevor analyzing the sample a background run was performed.

S2.4. Determination of the melting point

A small aliquot of mycosporine-cysteinolic acid was placed on a microscope slide and visually examined while the temperature was gradually increased from room temperature to 160 °C using a PolyTherm A Heiztischmikroskop from Wagner & Munz (Wagner & Munz GmbH, Munich, Germany).

S2.5. Stability during freeze-thaw cycles

An aqueous solution of mycosporine-cysteinolic acid (c = 2 mg/mL) was subjected to 10 freeze-thaw cycles. For each cycle, the solution was stored at -20 °C for 20 h and subsequently thawed at room temperature for 1 h. The sample was kept in standard glass HPLC vials with plastic screw caps.

### 3. Isolation trees

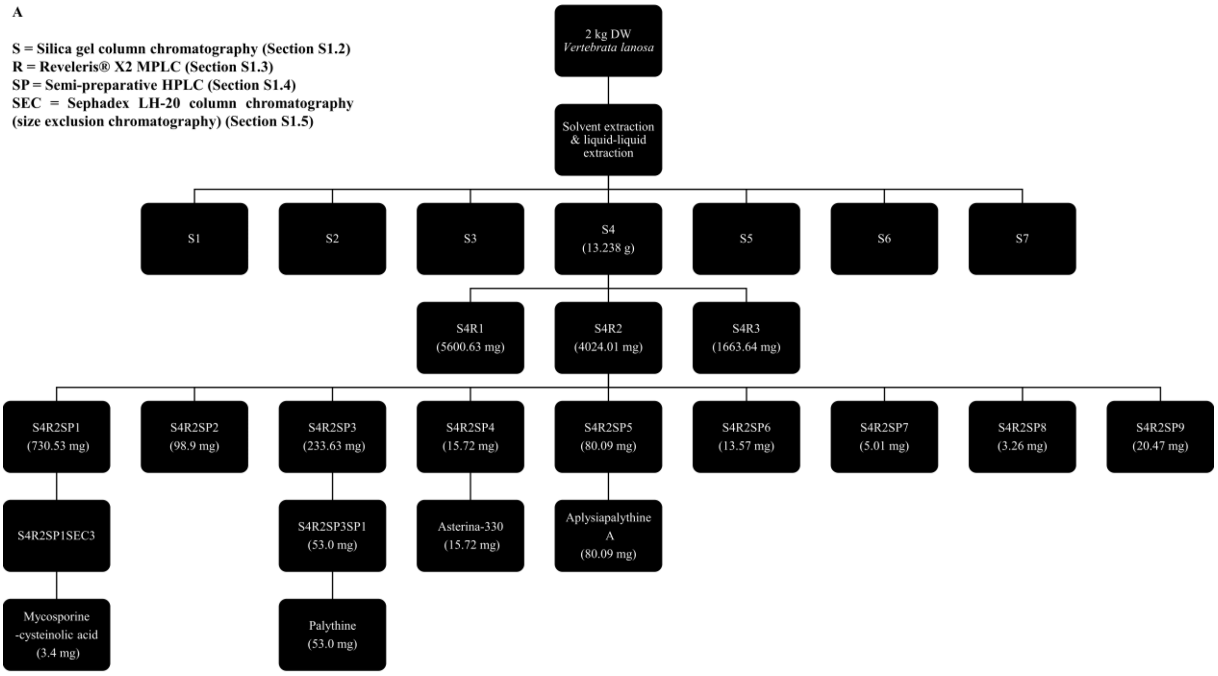

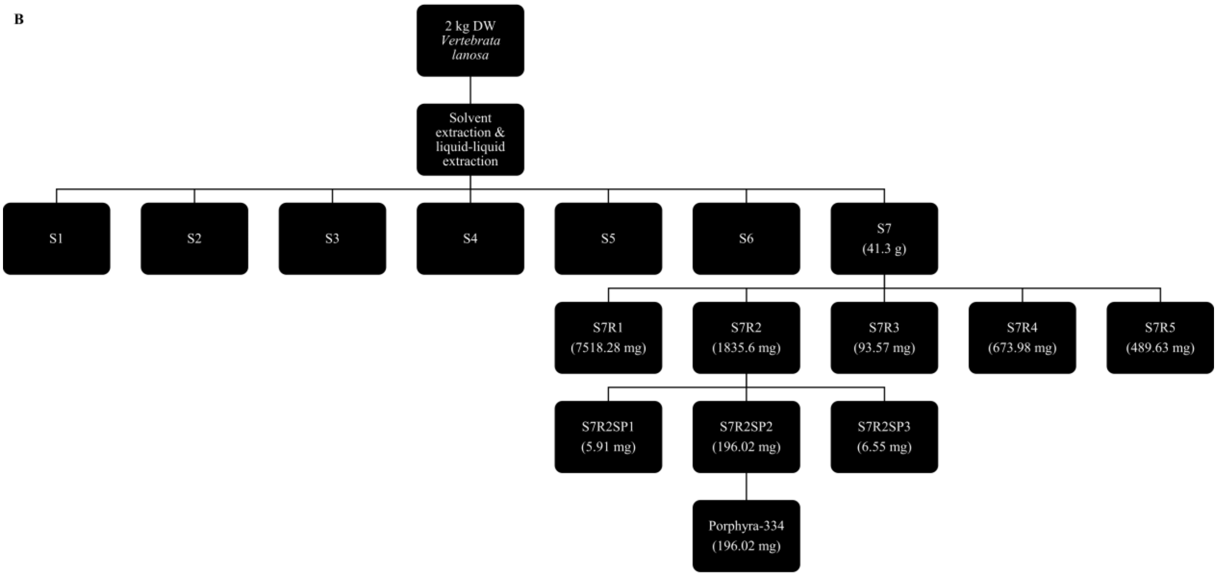

**Figure S1.** A | Isolation tree for fraction S4: 3.4 mg of the novel MAA mycosporine-cysteinolic acid, 53.0 mg of palythine, 15.72 mg of asterina-330, and 80.09 mg of aplysiapalythine A were obtained. B | Isolation tree for fraction S7: 196.02 mg of porphyra-334 were isolated.

#### 4. LC-MS analysis of mycosporine-cysteinolic acid

S4.1. HILIC-UPLC-DAD-MS analysis of fraction S4R2SP1SEC3 (mycosporine-cysteinolic acid)

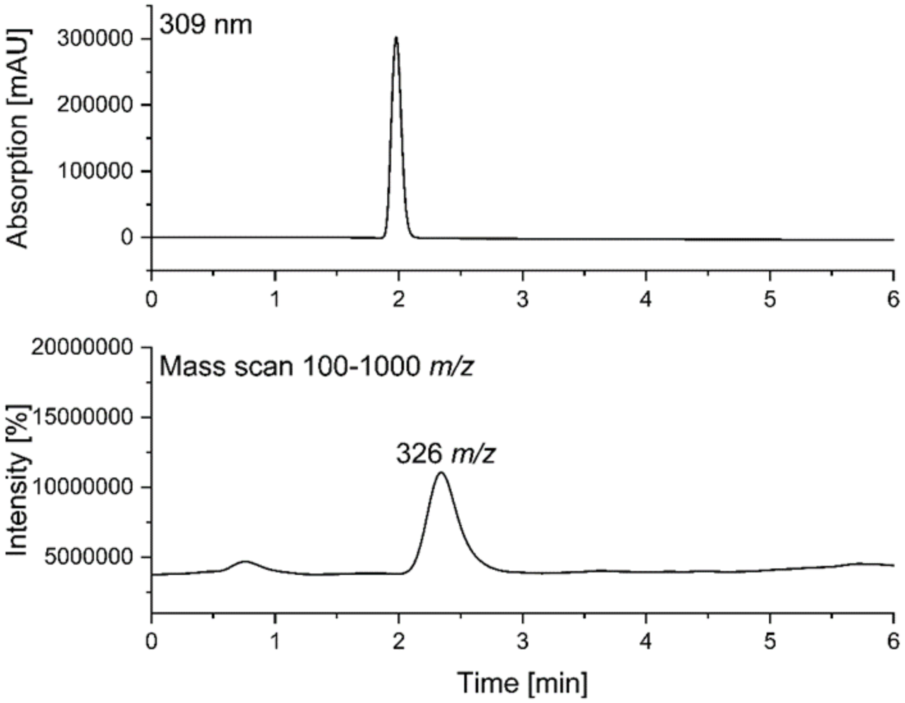

**Figure S2.** HILIC-UPLC-DAD-MS [1] of mycosporine-cysteinolic acid (S4R2SP1SEC3). Stationary phase: YMC Triart Diol HILIC column (50 × 2.1 mm, 1.9 µm particle size). Mobile phase: ACN/water (9:1, v/v) containing 5 mM ammonium acetate, adjusted to pH 6.5 with acetic acid (A), ACN/water (1:1, v/v) containing 15 mM ammonium acetate, adjusted to pH 3.5 with acetic acid (B). Gradient: in 10 min from 0% to 35% B; wash: for 10 min with 100% B; re-equilibration: for 10 min with initial conditions. Flow rate: 0.5 mL/min. Column oven temperature: 5 °C. Injection volume: 1 µL. MS conditions: ESI-positive-mode, capillary voltage 3.50 kV, cone voltage 50 V, collision energy 20 (eV), desolvation temperature 600 °C, and source temperature 150 °C.

#### 5. NMR data of isolated MAAs

S5.1. NMR spectra

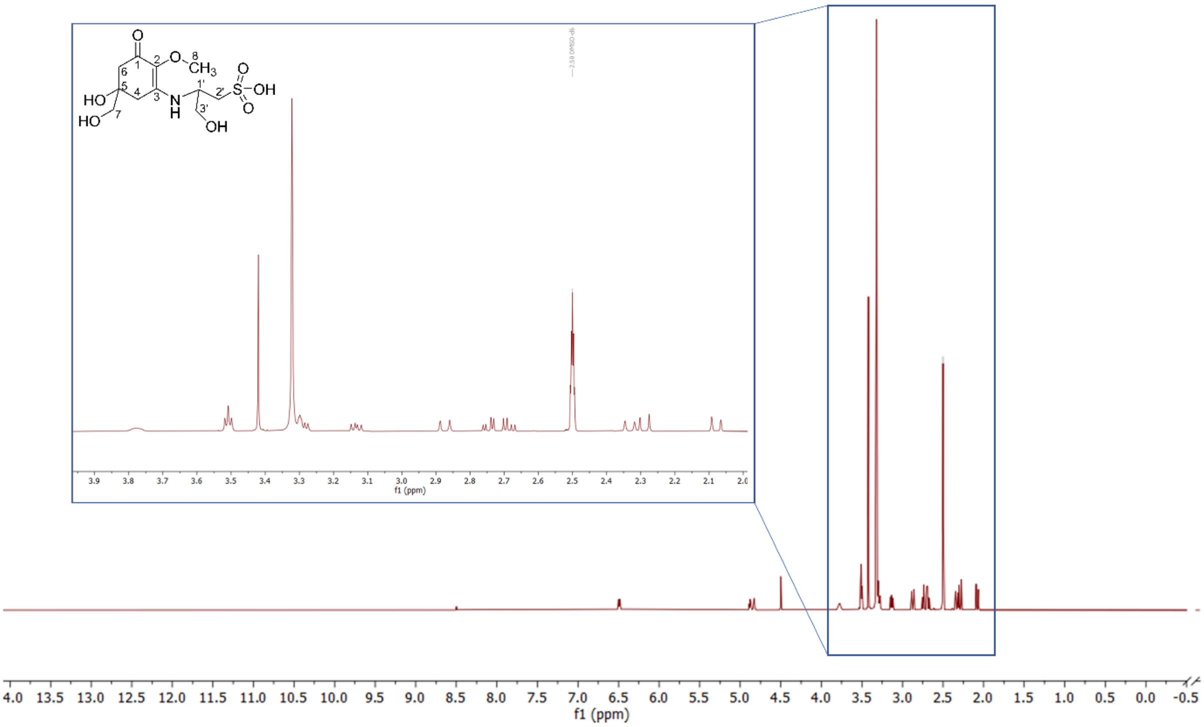

**Figure S3.** $^{1}$H NMR spectrum of the novel MAA mycosporine-cysteinolic acid (isolated from *Vertebrata lanosa*) recorded in DMSO-$d_6$ (600 MHz).

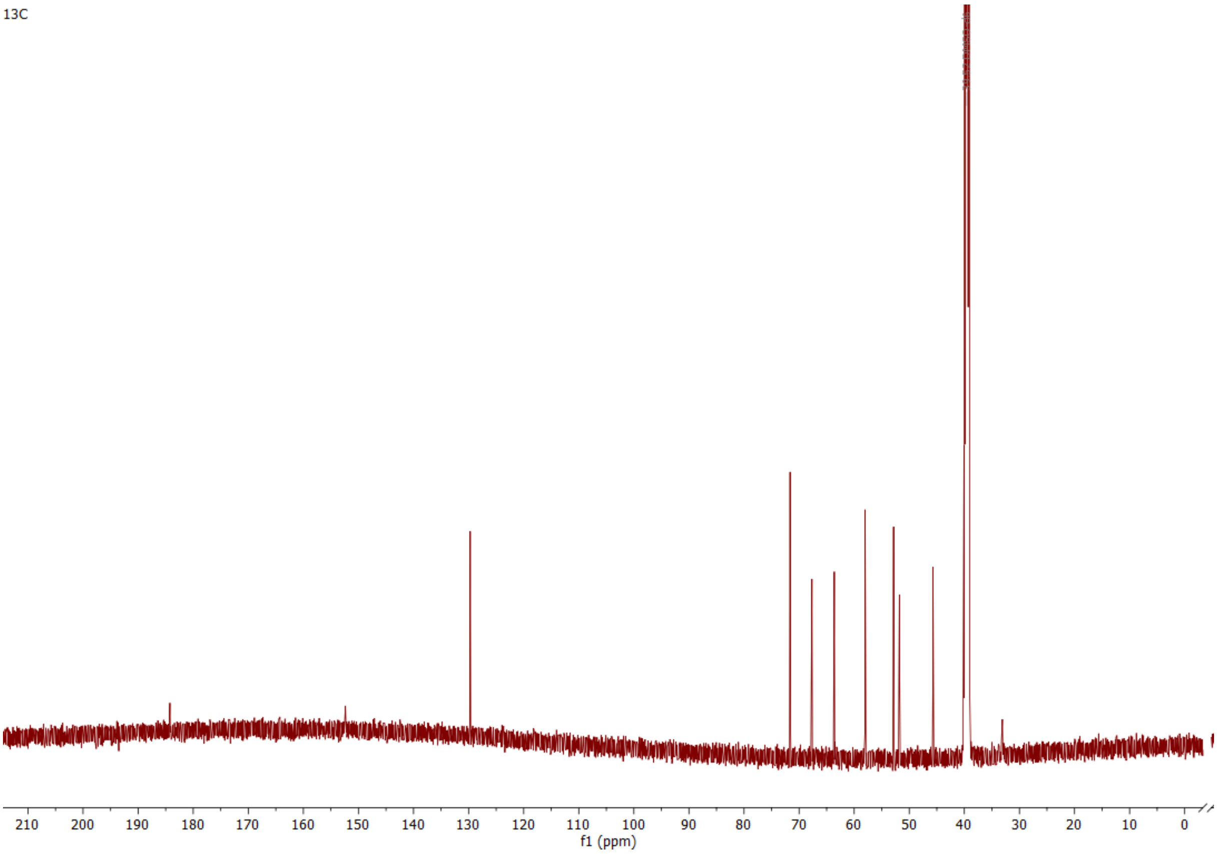

**Figure S4.** $^{13}C$ NMR spectrum of novel MAA (isolated from *Vertebrata lanosa*) recorded in DMSO-$d_6$ (151 MHz).

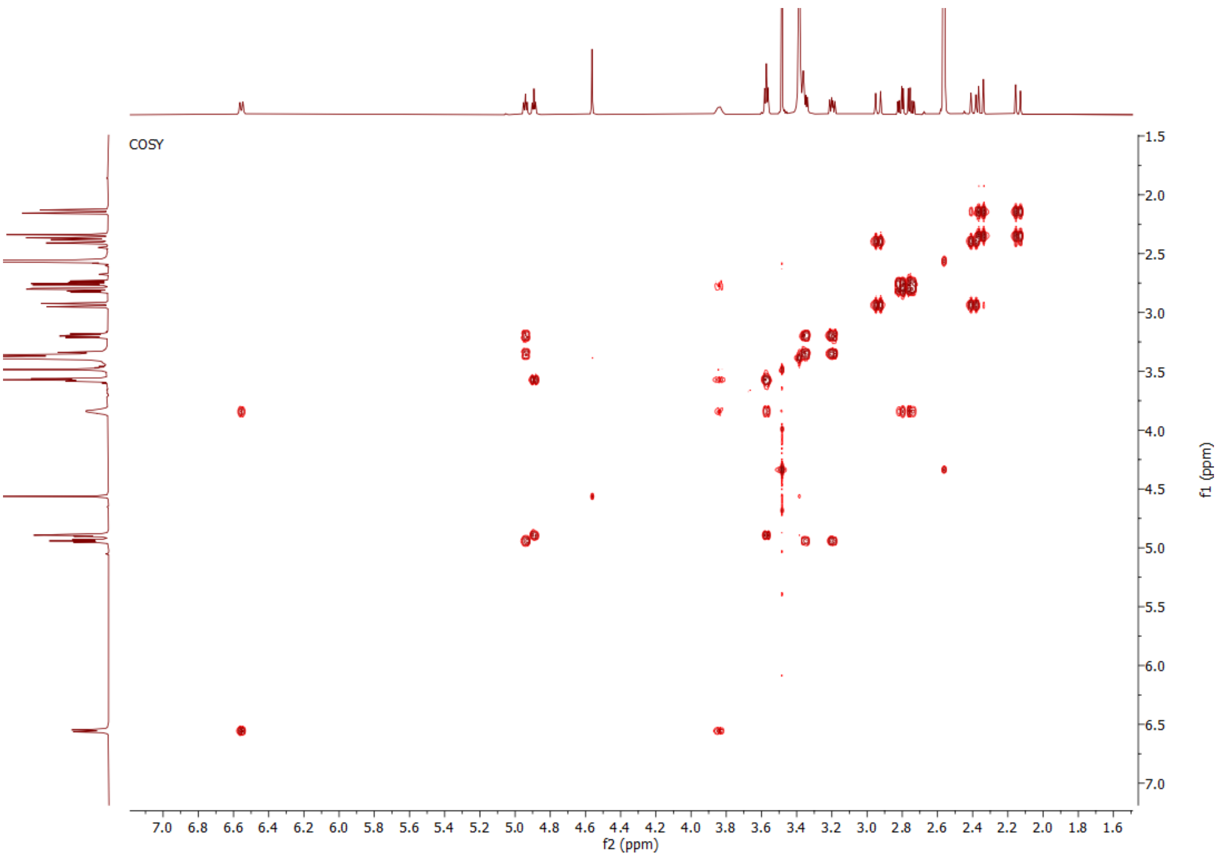

**Figure S5.** COSY NMR spectrum of mycosporine-cysteinolic acid recorded in DMSO-$d_6$ (600 MHz).

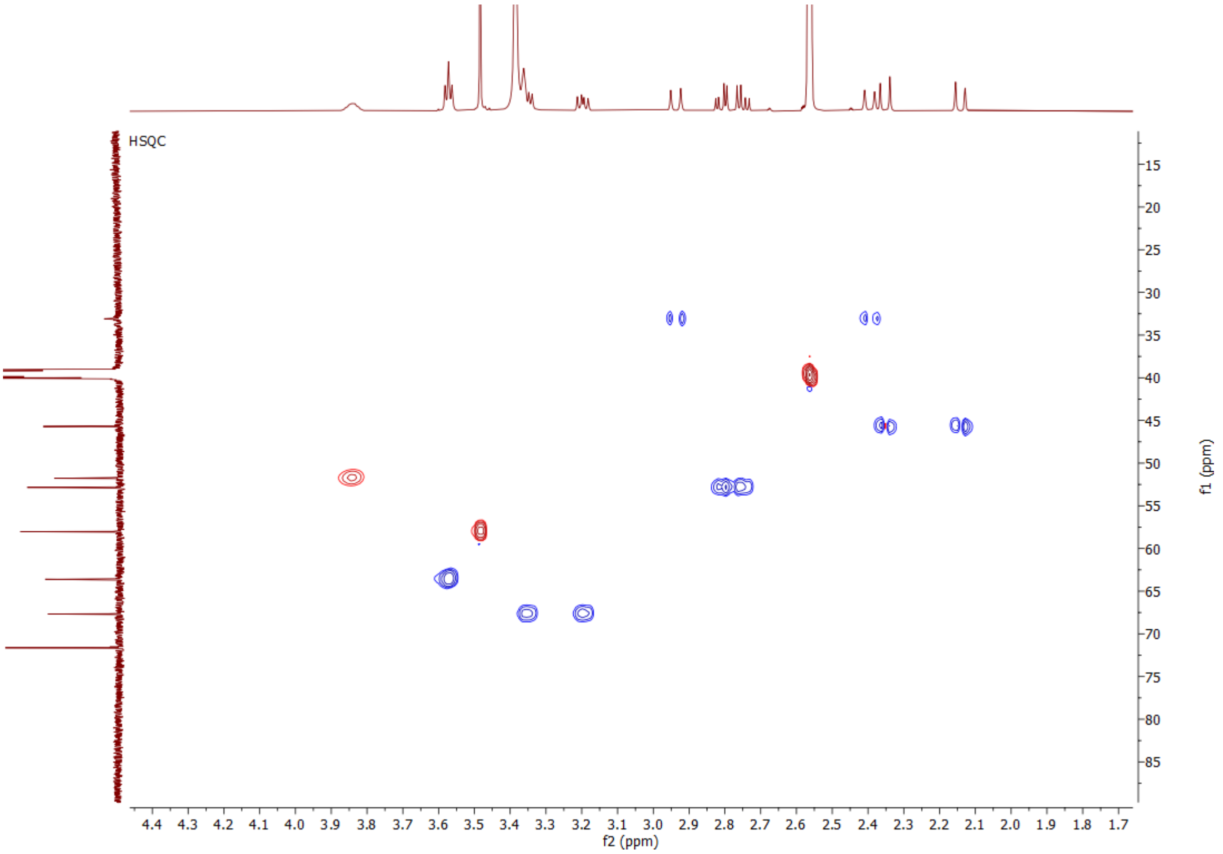

**Figure S6.** HSQC NMR spectrum of mycosporine-cysteinolic acid recorded in DMSO-$d_6$ (600 MHz, 151 MHz).

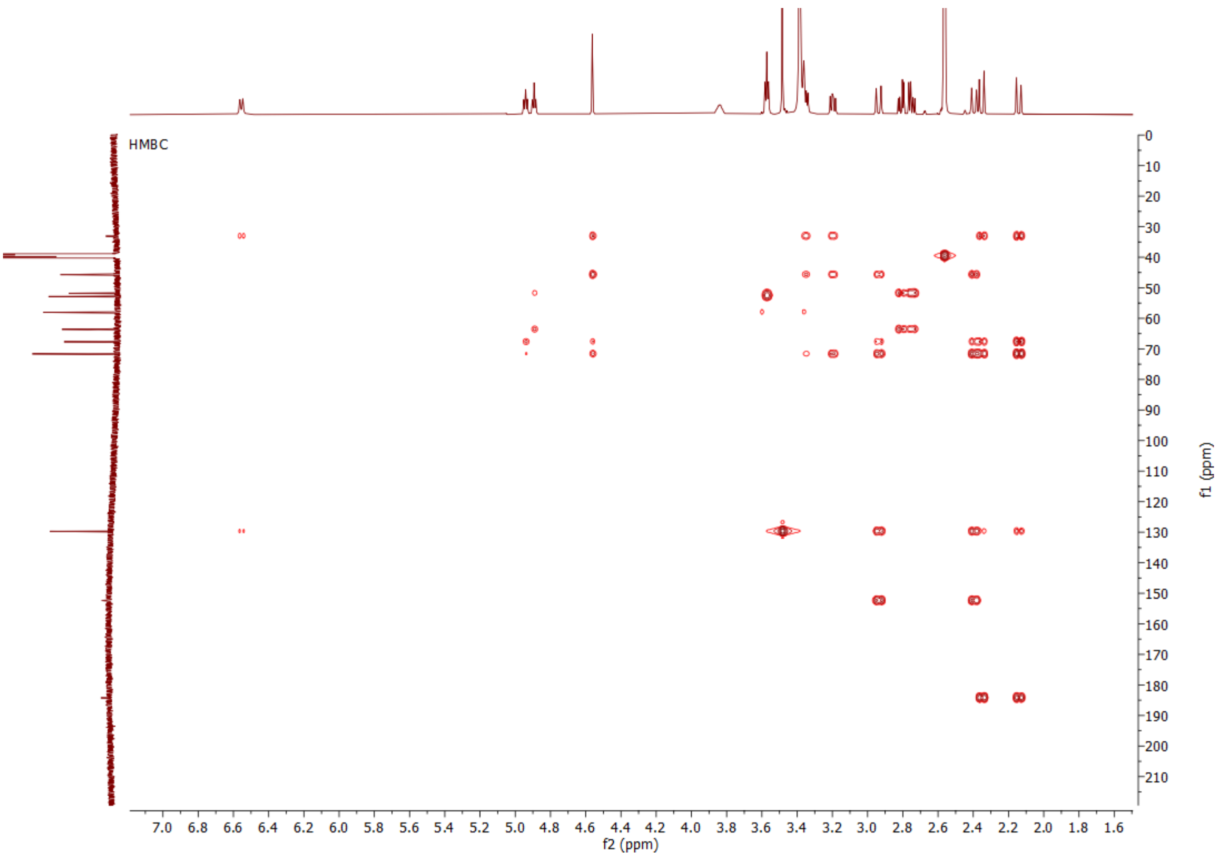

**Figure S7.** HMBC NMR spectrum of mycosporine-cysteinolic acid recorded in DMSO-$d_6$ (600 MHz, 151 MHz).

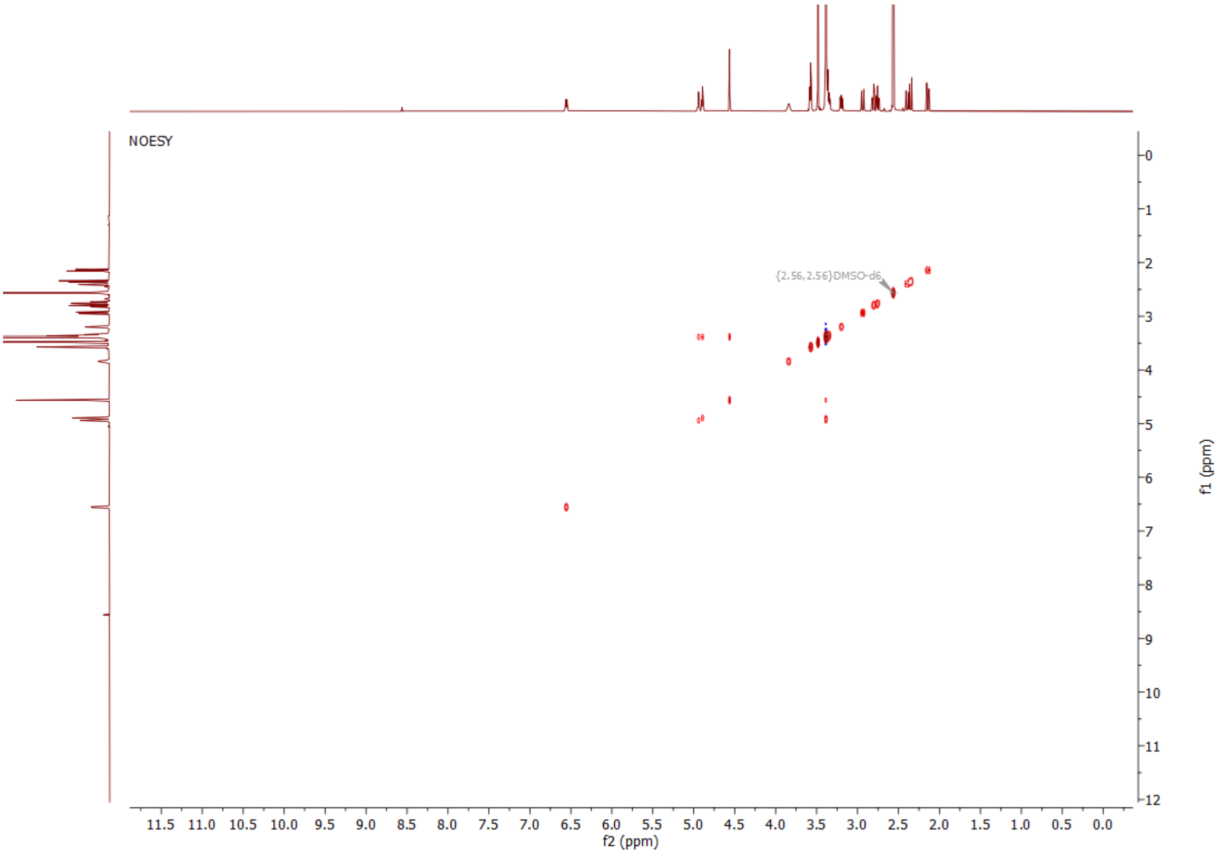

**Figure S8.** NOESY NMR spectrum of mycosporine-cysteinolic acid recorded in DMSO-$d_6$ (600 MHz).

S5.2. NMR data summary

NMR spectroscopic data of aplysiapalythine A: $^{1}$H NMR (400 MHz, DMSO-$d_6$) δ: 1.07 (d, $J$ = 6.17 Hz, 3H), 2.56 (d, $J$ = 17.26 Hz, 1H), 2.56 (d, $J$ = 17.26 Hz, 1H), 2.79 (d, $J$ = 11.05 Hz, 1H), 2.79 (d, $J$ = 11.05 Hz, 1H), 3.23 (dd, 2H), 3.28 (s, 2H), 3.49 (s, 3H), 3.64 (s, 2H), 3.77 (m, 1H), 8.27 (s, 1H), 8.39 (s, 1H). $^{13}$C NMR (101 MHz, DMSO-$d_6$) δ: 21.00, 33.68, 33.68, 50.28, 58.77, 65.61, 67.80, 70.67, 124.84, 159.03, 159.58, 163.89, 168.25.

NMR spectroscopic data of palythine: $^{1}$H NMR (400 MHz, DMSO-$d_6$) δ: 2.38 (d, $J$ = 17.01 Hz, 1H), 2.38 (d, $J$ = 17.01 Hz, 1H), 2.54 (d, $J$ = 10.31 Hz, 1H), 2.54 (d, $J$ = 10.31 Hz, 1H), 3.25 (s, 2H), 3.49 (s, 3H), 3.64 (s, 2H), 8.26 (s, 1H), 8.34 (s, 1H), 8.56 (s, 1H). $^{13}$C NMR (101 MHz, DMSO-$d_6$) δ: 34.32, 36.52, 46.90, 58.72, 67.71, 70.97, 124.31, 159.12, 161.29, 163.81, 168.00.

NMR spectroscopic data of porphyra-334: $^{1}$H NMR (400 MHz, DMSO-$d_6$) δ: 1.08 (d, $J$ = 5.76 Hz, 3H), 2.74 (2d, $J$ = 17.36 Hz, 2H), 2.83 (2d, $J$ = 16.93 Hz, 2H), 3.28 (s, 2H), 3.55 (s, 3H), 3.97 (d, 2H), 4.01 (d, 1H), 4.22 (dq, $J$ = 6.4, 4.6 Hz, 1H), 5.36–5.40 (s, 1H), 7.70 (s, 1H), 8.27 (s, 1H), 8.70 (s, 1H). $^{13}$C NMR (101 MHz, DMSO-$d_6$) δ: 20.56, 33.43, 33.93, 45.76, 59.13, 62.25, 67.29, 67.97, 125.35, 159.28, 160.54, 163.87, 170.02, 170.81.

NMR spectroscopic data of asterina-330: $^{1}H$ NMR (600 MHz, DMSO-$d_6$) δ: 2.56 (m, 2H), 2.78 (t, $J$ = 16.4 Hz, 2H), 3.28 (q, $J$ = 10.9 Hz, 4H), 3.49 (s, 3H), 3.53 (d, $J$ = 2.8 Hz, 2H), 3.63 (s, 2H), 8.29 (s, 1H), 8.38 (s, 1H). $^{13}C$ NMR (151 MHz, DMSO-$d_6$) δ: 33.50, 33.72, 45.53, 46.90, 58.75, 60.03, 67.82, 70.62, 124.85, 158.98, 159.40, 168.33.

Further experimental data are available upon request.

#### 6. IR spectrum of mycosporine-cysteinolic acid

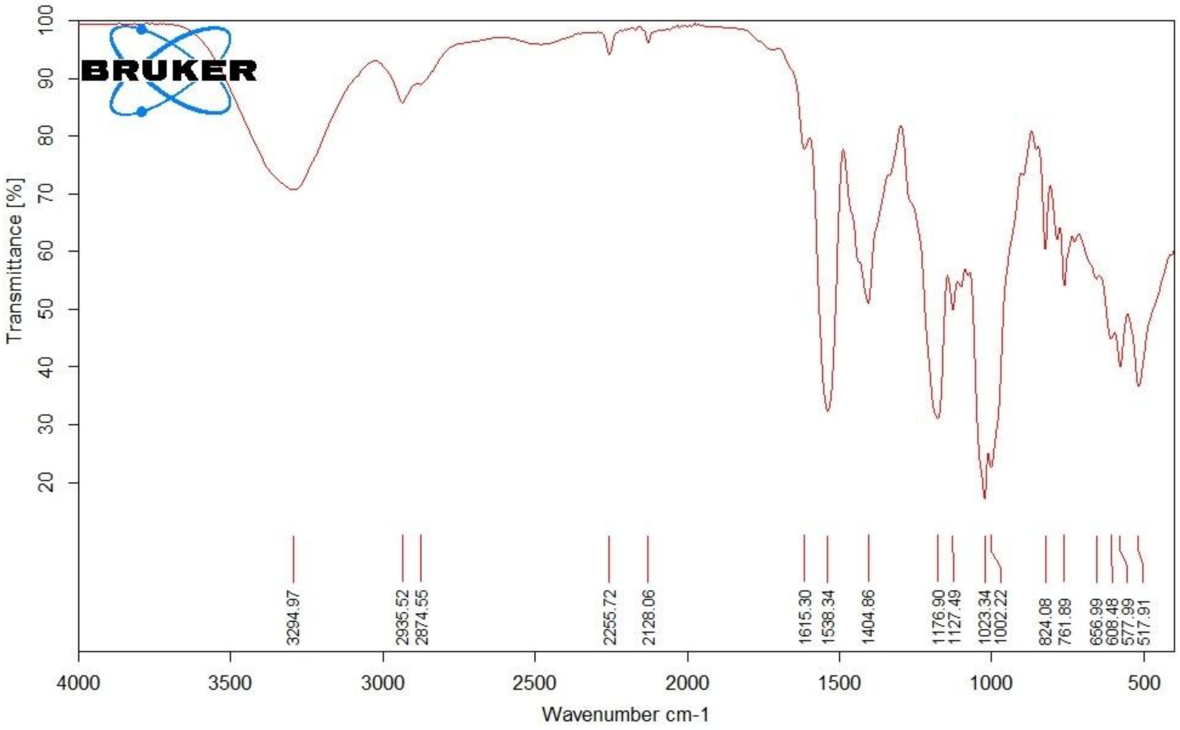

**Figure S9.** IR spectrum of mycosporine-cysteinolic acid.

#### 7. Circular dichroism spectrum of mycosporine-cysteinolic acid

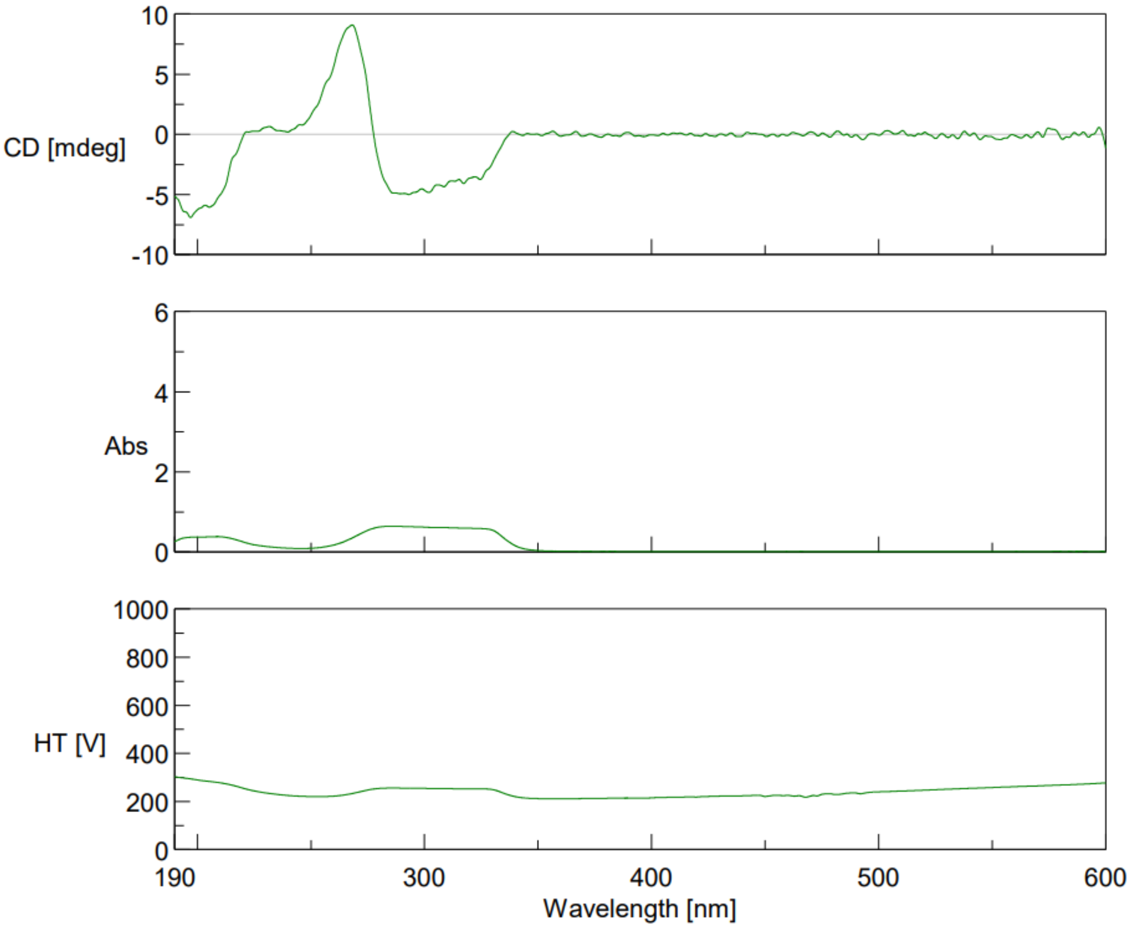

**Figure S10.** Circular dichroism spectrum of mycosporine-cysteinolic acid. CD ($H_2O$, 25 °C): $\lambda_{max}$ (mdeg) −6.8 (205 nm), +9.2 (285 nm), −5.0 (310 nm) (c = 210 μg/mL).

#### 8. Calculated electronic circular dichroism (ECD) spectra

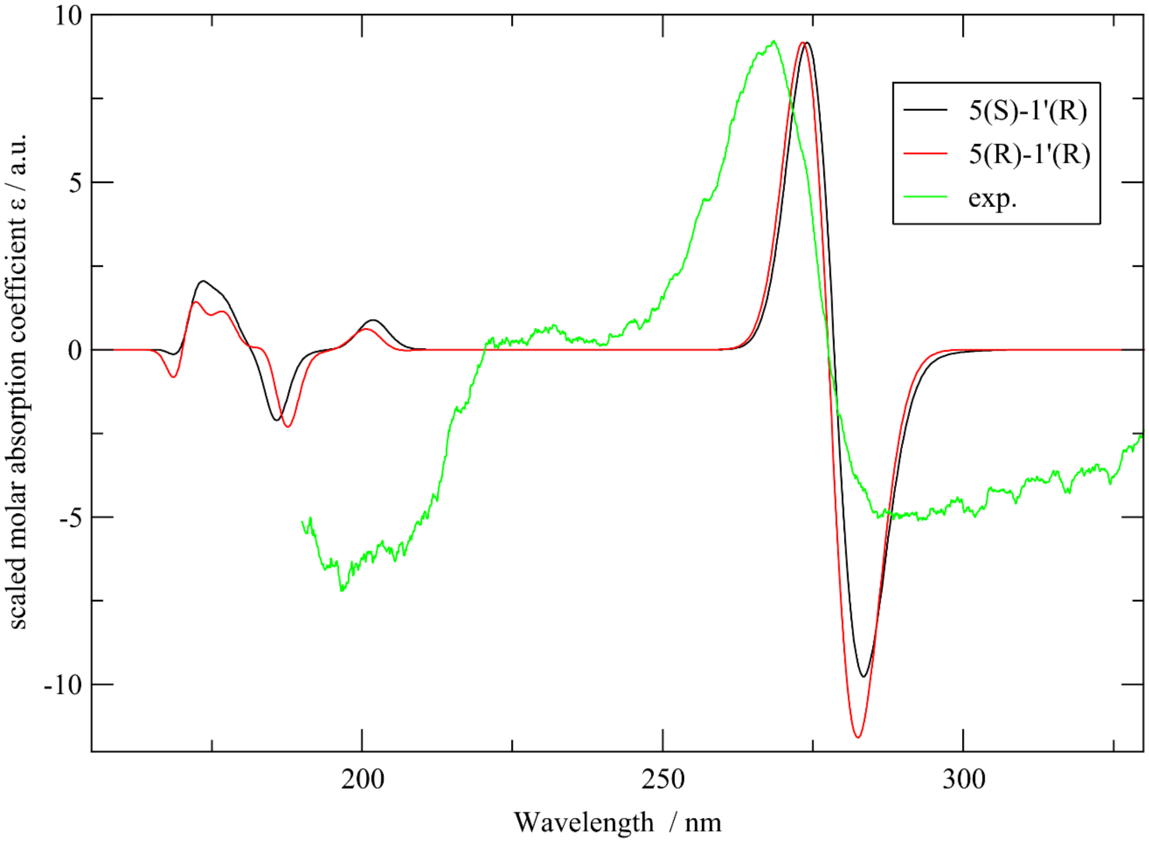

**Figure S11.** The experimental CD spectrum of the isolated MAA (mycosporine-cysteinolic acid) as well as computed ECD spectra of the SR and RR diastereomers.

**Table S1**. Conformationally averaged NMR chemical shifts of the carbon atoms in RR and SR stereoisomers. All reported values are in ppm. Exp.: experimental values, original: average chemical shifts as directly obtained from DFT, corrected: DFT original values are offset by the average error, error: difference between corrected and experimental value. The errors marked in red are those with an absolute value above 2 ppm.

**Carbon:**

| Atom # | Exp. | 5R-1'R original | 5R-1'R corrected | 5R-1'R error | 5S-1'R original | 5S-1'R corrected | 5S-1'R error |
|---|---|---|---|---|---|---|---|
| 1 | 184.20 | 191.11 | 183.96 | -0.24 | 191.93 | 184.32 | 0.12 |
| 2 | 129.70 | 137.80 | 130.65 | 0.95 | 138.10 | 130.48 | 0.78 |
| 3 | 152.40 | 161.36 | 154.22 | 1.82 | 161.41 | 153.80 | 1.40 |
| 4 | 33.10 | 38.84 | 31.69 | -1.41 | 37.87 | 30.26 | **-2.84** |
| 5 | 71.60 | 80.43 | 73.28 | 1.68 | 80.70 | 73.09 | 1.49 |
| 6 | 45.70 | 52.04 | 44.89 | -0.81 | 51.14 | 43.53 | **-2.17** |
| 7 | 67.70 | 73.43 | 66.28 | -1.42 | 78.59 | 70.98 | **3.28** |
| 8 | 58.00 | 62.28 | 55.13 | **-2.87** | 62.56 | 54.94 | **-3.06** |
| 1' | 51.80 | 60.31 | 53.16 | 1.36 | 60.23 | 52.61 | 0.81 |
| 2' | 52.90 | 57.39 | 50.24 | **-2.66** | 57.80 | 50.19 | **-2.71** |
| 3' | 63.60 | 74.35 | 67.20 | **3.60** | 74.10 | 66.49 | **2.89** |
| RMSD (total) | | | | 1.95 | | | **2.22** |
| RMSD (4 – 7) | | | | 1.37 | | | **2.54** |

#### 9. Analyzed algal samples

**Table S2**. Information on the analyzed algae.

| Species | Family, Order | Collection Place | Country | Sampling season | Life stage | Code | Collection Date |
|---|---|---|---|---|---|---|---|
| *Porphyra* sp. commercial sample | Bangiaceae, Bangiales | Not known* | China | Not known* | Not known* | - | Not known* |
| *Vertebrata lanosa* (I) | Rhodomelaceae, Ceramiales | Portsall | France | Summer | Not known* | - | 8/2024 |
| *Vertebrata lanosa* (II) | Rhodomelaceae, Ceramiales | Roscoff | France | Spring | Not known* | - | 6/2018 |
| *Cryptopleura cryptoneuron* | Delesseriaceae, Ceramiales | Paracas District | Peru | Summer | Gametophyte | 1.3 | 3/24 |
| *Griffithsia pacifica* | Wrangeliaceae, Ceramiales | Paracas District | Peru | Summer | Gametophyte | 2.1 | 3/24 |
| *Tiffaniella snyderae* | Wrangeliaceae, Ceramiales | Paracas District | Peru | Summer | Gametophyte | 3.1 | 3/24 |
| *Corallina chilensis* | Corallinaceae, Corallinales | Paracas District | Peru | Spring | Gametophyte | 4.1 | 11/23 |
| *Asterfilopsis centralis* | Phyllophoraceae, Gigartinales | Paracas District | Peru | Spring | Gametophyte | 5.1 | 11/23 |
| *Sarcodiotheca gaudichaudii* | Soleriaceae, Gigartinales | Paracas District | Peru | Spring | Gametophyte | 6.1 | 11/23 |

| | | | | | | | |
|---|---|---|---|---|---|---|---|
| *Iridaea tuberculosa* | Gigartinaceae, Gigartinales | Paracas District | Peru | Spring | Gametophyte | 7.1 | 11/23 |
| *Chondrus canaliculatus* | Gigartinaceae, Gigartinales | Paracas District | Peru | Spring | Gametophyte | 8.1 | 11/23 |
| *Chondracanthus chamissoi* | Gigartinaceae, Gigartinales | Paracas District | Peru | Spring | Gametophyte | 9.1 | 11/23 |
| *Nitophyllum peruvianum* | Delesseriaceae, Ceramiales | Paracas District | Peru | Summer | Gametophyte | 10.1 | 3/24 |
| *Gracilariopsis lemaneiformis* | Gracilariaceae, Gracilariales | Paracas District | Peru | Spring | Gametophyte | 11.1 | 11/23 |
| *Prionitis decipiens* | Halymeniaceae, Halymeniales | Paracas District | Peru | Spring | Sporophyte | 12.1 | 11/23 |
| *Rhodymenia flabellifolia* | Rhodymeniaceae, Rhodymeniales | Paracas District | Peru | Summer | Gametophyte | 13.1 | 3/24 |

#### 10.Feature-based molecular networking

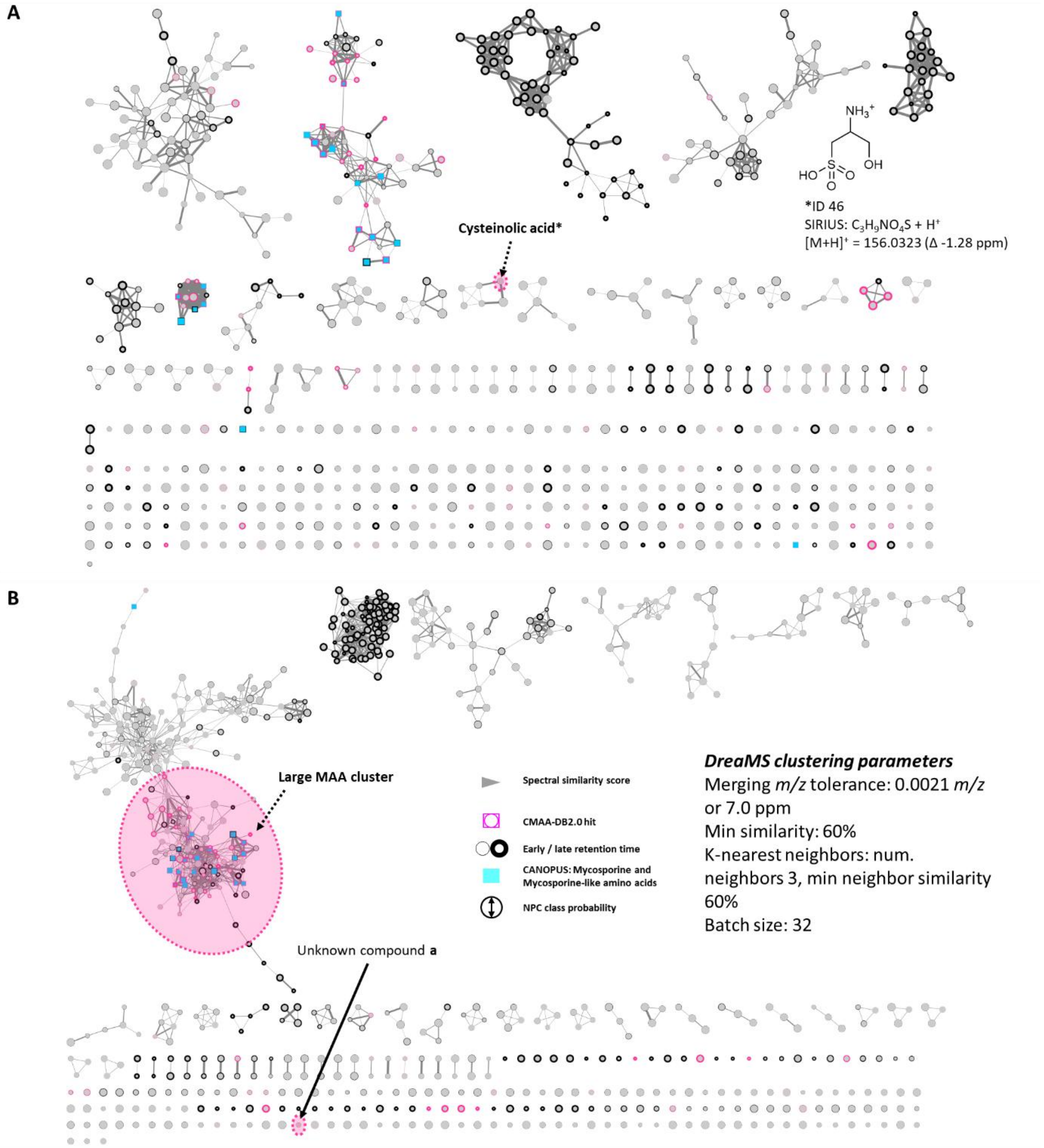

**Figure S12.** A | FBMN based on the cosine similarity score, with the feature (ID 46) putatively corresponding to cysteinolic acid highlighted. B | FBMN using the DreaMS similiarity scores for clustering.

#### 11.CMAA-DB2.0

**Table S3.** Summary of the SMILES input strings used to generate the new combinatorial MAA database (CMAA-DB2.0) including compound class identifications with ClassyFire [2] (see http://classyfire.wishartlab.com/queries/12793695) and biological sources of (some) listed compounds.

| # | Name | Compound class (ClassyFire subclass level) | Occurring in … | Organism(s) | Complete SMILES | SmiLib building block | Type | Reference |
|---|---|---|---|---|---|---|---|---|
| | Cyclohexenone-type | | | | | O=C1C(OC)=C(CC(O)(C1)CO)N[R1] | **Scaffold** | |
| | Algasporine-type | | | | | O=C1C(OC)=C(N([R1])C)CC(C1O)(CO)O | **Scaffold** | |
| | Deoxyalgasporine-type | | | | | O=C1C(OC)=C(CC(CO)(O)C1)N(C)[R1] | **Scaffold** | |
| | Hydroxycyclohexenone-type | | | | | O=C1C(OC)=C(CC(O)(C1O)CO)N[R1] | **Scaffold** | |
| | Palythine-type | | | | | COC1=C(CC(O)(CC1=N)CO)N[R1] | **Scaffold** | |
| | N-Methylhydroxypalythine-type | | | | | COC1=C(CC(O)(C(O)C1=N)CO)N(C)[R1] | **Scaffold** | |
| | N-Methylpalythine-type | | | | | COC1=C(CC(O)(CC1=N)CO)N(C)[R1] | **Scaffold** | |
| | Hydroxypalythine-type | | | | | COC1=C(CC(O)(C(O)C1=N)CO)N[R1] | **Scaffold** | |
| | Aminocyclohexenimine-type | | | | | COC1=C(CC(O)(CC1=N[R2])CO)N[R1] | **Scaffold** | |
| | N-Methylhydroxy-aminocyclohexenimine-type | | | | | COC1=C(CC(O)(C(O)C1=N[R2])CO)N(C)[R1] | **Scaffold** | |
| | N-Methylamino-cyclohexenimine-type | | | | | COC1=C(CC(O)(CC1=N[R2])CO)N(C)[R1] | **Scaffold** | |
| | Hydroxyamino-cyclohexenimine-type | | | | | COC1=C(CC(O)(C(O)C1=N[R2])CO)N[R1] | **Scaffold** | |
| | | | | | | [A][R1] | **Linker** | |

| | | | | | | | |
|---|---|---|---|---|---|---|---|
| 1 | Arginine | Amino acids, peptides, and analogues | | | NC(CCCNC(N)=N)C(O)=O | [A]C(CCCNC(N)=N)C(O)=O | **Building block** |
| 2 | Arginine-$CO_2$ | Guanidines | | | NCCCCNC(N)=N | [A]CCCCNC(N)=N | **Building block** |
| 3 | Histidine | Amino acids, peptides, and analogues | | | NC(CC1=CNC=N1)C(O)=O | [A]C(CC1=CNC=N1)C(O)=O | **Building block** |
| 4 | Histidine-$CO_2$ | Amines | | | NCCC1=CNC=N1 | [A]CCC1=CNC=N1 | **Building block** |
| 5 | Lysine | Amino acids, peptides, and analogues | | | NC(CCCCN)C(O)=O | [A]C(CCCCN)C(O)=O | **Building block** |
| 6 | Lysine-$CO_2$ | Amines | | | NCCCCCN | [A]CCCCCN | **Building block** |
| 7 | Aspartic acid | Amino acids, peptides, and analogues | | | NC(CC(O)=O)C(O)=O | [A]C(CC(O)=O)C(O)=O | **Building block** |
| 8 | Aspartic acid-$CO_2$ | Amino acids, peptides, and analogues | | | NCCC(O)=O | [A]CCC(O)=O | **Building block** |
| 9 | Glutamic acid | Amino acids, peptides, and analogues | | | NC(CCC(O)=O)C(O)=O | [A]C(CCC(O)=O)C(O)=O | **Building block** |
| 10 | Glutamic acid-$CO_2$ | Amino acids, peptides, and analogues | | | NCCCC(O)=O | [A]CCCC(O)=O | **Building block** |
| 11 | Serine | Amino acids, peptides, and analogues | | | NC(CO)C(O)=O | [A]C(CO)C(O)=O | **Building block** |
| 12 | Serine-$CO_2$ | Amines | | | NCCO | [A]CCO | **Building block** |
| 13 | Threonine | Amino acids, peptides, and analogues | | | NC(C(O)C)C(O)=O | [A]C(C(O)C)C(O)=O | **Building block** |
| 14 | Threonine-$CO_2$ = Threamine | Amines | | | NCC(O)C | [A]CC(O)C | **Building block** |
| 15 | Asparagine | Amino acids, peptides, and analogues | | | NC(CC(N)=O)C(O)=O | [A]C(CC(N)=O)C(O)=O | **Building block** |
| 16 | Asparagine-$CO_2$ | Amino acids, peptides, and analogues | | | NCCC(N)=O | [A]CCC(N)=O | **Building block** |

| | | | | | | | |
|---|---|---|---|---|---|---|---|
| 17 | Glutamine | Amino acids, peptides, and analogues | | | NC(CCC(N)=O)C(O)=O | [A]C(CCC(N)=O)C(O)=O | **Building block** |
| 18 | Glutamine-$CO_2$ | Amino acids, peptides, and analogues | | | NCCCC(N)=O | [A]CCCC(N)=O | **Building block** |
| 19 | Cysteine | Amino acids, peptides, and analogues | | | NC(CS)C(O)=O | [A]C(CS)C(O)=O | **Building block** |
| 20 | Cysteine-$CO_2$ | Alkylthiols | | | NCCS | [A]CCS | **Building block** |
| 21 | Selenocysteine | Amino acids, peptides, and analogues | | | NC(C[SeH])C(O)=O | [A]C(C[SeH])C(O)=O | **Building block** |
| 22 | Selenocysteine-$CO_2$ | Organoselenium compounds | | | NCC[SeH] | [A]CC[SeH] | **Building block** |
| 23 | Glycine | Amino acids, peptides, and analogues | | | NCC(O)=O | [A]CC(O)=O | **Building block** |
| 24 | Glycine-$CO_2$ | Amines | | | NC | [A]C | **Building block** |
| 25 | Alanine | Amino acids, peptides, and analogues | | | NC(C)C(O)=O | [A]C(C)C(O)=O | **Building block** |
| 26 | Alanine-$CO_2$ | Amines | | | NCC | [A]CC | **Building block** |
| 27 | Isoleucine | Amino acids, peptides, and analogues | | | NC(C(C)CC)C(O)=O | [A]C(C(C)CC)C(O)=O | **Building block** |
| 28 | Isoleucine-$CO_2$ | Amines | | | NCC(C)CC | [A]CC(C)CC | **Building block** |
| 29 | Leucine | Amino acids, peptides, and analogues | | | NC(CC(C)C)C(O)=O | [A]C(CC(C)C)C(O)=O | **Building block** |
| 30 | Leucine-$CO_2$ | Amines | | | NCCC(C)C | [A]CCC(C)C | **Building block** |
| 31 | Methionine | Amino acids, peptides, and analogues | | | NC(CCSC)C(O)=O | [A]C(CCSC)C(O)=O | **Building block** |
| 32 | Methionine-$CO_2$ | Dialkylthioethers | | | NCCCSC | [A]CCCSC | **Building block** |

| | | | | | | | |
|---|---|---|---|---|---|---|---|
| 33 | Phenylalanine | Amino acids, peptides, and analogues | | | NC(CC1=CC=CC=C1)C(O)=O | [A]C(CC1=CC=CC=C1)C(O)=O | **Building block** |
| 34 | Phenylalanine-$CO_2$ | Phenethylamines | | | NCCC1=CC=CC=C1 | [A]CCC1=CC=CC=C1 | **Building block** |
| 35 | Tryptophan | Indolyl carboxylic acids and derivatives | | | NC(CC1=CNC2=C1C=CC=C2)C(O)=O | [A]C(CC1=CNC2=C1C=CC=C2)C(O)=O | **Building block** |
| 36 | Tryptophan-$CO_2$ | Tryptamines and derivatives | | | NCCC1=CNC2=C1C=CC=C2 | [A]CCC1=CNC2=C1C=CC=C2 | **Building block** |
| 37 | Valine | Amino acids, peptides, and analogues | | | NC(C(C)C)C(O)=O | [A]C(C(C)C)C(O)=O | **Building block** |
| 38 | Valine-$CO_2$ | Amines | | | NCC(C)C | [A]CC(C)C | **Building block** |
| 39 | Tyrosine | Amino acids, peptides, and analogues | | | NC(CC1=CC=C(O)C=C1)C(O)=O | [A]C(CC1=CC=C(O)C=C1)C(O)=O | **Building block** |
| 40 | Tyrosine-$CO_2$ | Phenethylamines | | | NCCC1=CC=C(O)C=C1 | [A]CCC1=CC=C(O)C=C1 | **Building block** |
| 41 | Prop-1-en-1-amine | Amines | | | NC=CC | [A]C=CC | **Building block** |
| 42 | Taurine | Organosulfonic acids and derivatives | | | OS(=O)(CCN)=O | OS(=O)(CC[A])=O | **Building block** |
| 43 | Ornithine | Amino acids, peptides, and analogues | | | NC(CCCN)C(O)=O | [A]C(CCCN)C(O)=O | **Building block** |
| 44 | Ornithine-$CO_2$ | Amines | | | NCCCCN | [A]CCCCN | **Building block** |
| 45 | Citrulline | Amino acids, peptides, and analogues | | | NC(CCCNC(N)=O)C(O)=O | [A]C(CCCNC(N)=O)C(O)=O | **Building block** |
| 46 | Citrulline-$CO_2$ | Carboximidic acids | | | NCCCCNC(N)=O | [A]CCCCNC(N)=O | **Building block** |
| 47 | DOPA | Amino acids, peptides, and analogues | | | OC(C(CC1=CC=C(O)C(O)=C1)N)=O | OC(C(CC1=CC=C(O)C(O)=C1)[A])=O | **Building block** |
| 48 | DOPA-$CO_2$ | Benzenediols | | | NCCC1=CC=C(O)C(O)=C1 | [A]CCC1=CC=C(O)C(O)=C1 | **Building block** |

| | | | | | | | |
|---|---|---|---|---|---|---|---|
| 49 | 5-Hydroxytryptophan | Tryptamines and derivatives | | | NC(CC1=CNC2=C1C=C(C=C2)O)C(O)=O | [A]C(CC1=CNC2=C1C=C(C=C2)O)C(O)=O | **Building block** |
| 50 | 5-Hydroxytryptophan-$CO_2$ | Tryptamines and derivatives | | | NCCC1=CNC2=C1C=C(C=C2)O | [A]CCC1=CNC2=C1C=C(C=C2)O | **Building block** |
| 51 | β-N-Methylaminoalanine | Amino acids, peptides, and analogues | | | NC(C(O)=O)CNC | [A]C(C(O)=O)CNC | **Building block** |
| 52 | β-N-Methylaminoalanine-$CO_2$ | Amines | | | NCCNC | [A]CCNC | **Building block** |
| 53 | Ibotenic acid | Amino acids, peptides, and analogues | | | NC(C1=CC(NO1)=O)C(O)=O | [A]C(C1=CC(NO1)=O)C(O)=O | **Building block** |
| 54 | Ibotenic acid-$CO_2$ | Amines | | | NCC1=CC(NO1)=O | [A]CC1=CC(NO1)=O | **Building block** |
| 55 | Argininosuccinic acid | Amino acids, peptides, and analogues | | | O=C(O)C(N)CCCN=C(NC(C(O)=O)CC(O)=O)N | O=C(O)C([A])CCCN=C(NC(C(O)=O)CC(O)=O)N | **Building block** |
| 56 | Argininosuccinic acid-$CO_2$ | Amino acids, peptides, and analogues | | | NCCCCN=C(NC(C(O)=O)CC(O)=O)N | [A]CCCCN=C(NC(C(O)=O)CC(O)=O)N | **Building block** |
| 57 | Homoserine | Amino acids, peptides, and analogues | | | NC(CCO)C(O)=O | [A]C(CCO)C(O)=O | **Building block** |
| 58 | Homoserine-$CO_2$ | Amines | | | NCCCO | [A]CCCO | **Building block** |
| 59 | 3-Aminoisobutyric acid | Amino acids, peptides, and analogues | | | NCC(C)C(O)=O | [A]CC(C)C(O)=O | **Building block** |
| 60 | 3-Aminoisobutyric acid-$CO_2$ | Amines | | | NCCC | [A]CCC | **Building block** |
| 61 | p-Aminobenzoic acid | Benzoic acids and derivatives | | | O=C(C1=CC=C(C=C1)N)O | O=C(C1=CC=C(C=C1)[A])O | **Building block** |
| 62 | p-Aminobenzoic acid-$CO_2$ | Aniline and substituted anilines | | | NC1=CC=CC=C1 | [A]C1=CC=CC=C1 | **Building block** |
| 63 | Dehydroalanine | Amino acids, peptides, and analogues | | | NC(C(O)=O)=C | [A]C(C(O)=O)=C | **Building block** |

| | | | | | | | | |
|---|---|---|---|---|---|---|---|---|
| 64 | Dehydroalanine-$CO_2$ | Amines | | | NC=C | [A]C=C | **Building block** | |
| 65 | Homocysteine | Amino acids, peptides, and analogues | | | NC(CCS)C(O)=O | [A]C(CCS)C(O)=O | **Building block** | |
| 66 | Homocysteine-$CO_2$ | Alkylthiols | | | NCCCS | [A]CCCS | **Building block** | |
| 67 | Phosphoserine | Amino acids, peptides, and analogues | Red algae, brown algae | *Porphyra* sp., *Undaria pinnatifia*, *Laminaria* sp., *Sargassum fusiforme*, *Saccharina latissima*, *Laminaria digitata*, *Fucus vesiculosus*, *Ascophyllum nodosum* | NC(COP(O)(O)=O)C(O)=O | [A]C(COP(O)(O)=O)C(O)=O | **Building block** | [3, 4] |
| 68 | Phosphoserine-$CO_2$ | Phosphate esters | | | NCCOP(O)(O)=O | [A]CCOP(O)(O)=O | **Building block** | |
| 69 | Cystathionine | Amino acids, peptides, and analogues | Red algae, brown algae, green microalgae | *Saccharina latissima*, *Laminaria digitata*, *Fucus vesiculosus*, *Ascophyllum nodosum*, *Devaleraea* sp., *Pyropia pseudolinearis*, … | O=C(O)C(N)CSCCC(N)C(O)=O | O=C(O)C(N)CSCCC([A])C(O)=O<br>O=C(O)C([A])CSCCC(N)C(O)=O | **Building block** | [4, 5] |

| | | | | | | | | |
|---|---|---|---|---|---|---|---|---|
| 70 | Cystathionine-$CO_2$ | Amino acids, peptides, and analogues | | | O=C(O)C(N)CSCCCN NCCSCCC(N)C(O)=O | O=C(O)C(N)CSCCC[A] [A]CCSCCC(N)C(O)=O | **Building block** | |
| 71 | Phosphoethanolamine | Phosphate esters | Red algae, brown algae | *Porphyra* sp., *Undaria pinnatifia*, *Laminaria* sp., *Sargassum fusiforme* | OP(O)(OCCN)=O | OP(O)(OCC[A])=O | **Building block** | [3] |
| 72 | Hypotaurine | Sulfinic acids and derivatives | Marine invertebrates (snail in hydrothermal vents, hydrothermal vent tubeworm) | *Depressigyra globulus*, *Riftia* sp. | OS(CCN)=O | OS(CC[A])=O | **Building block** | [6] |
| 73 | Thiotaurine | Thiosulfonic acid | Marine invertebrates (snail in hydrothermal vents) | *Depressigyra globulus* | S=S(O)(CCN)=O | S=S(O)(CC[A])=O | **Building block** | [6] |
| 74 | Proline | Amino acids, peptides, and analogues | Microalgae | *Scenedesmus* sp. | O=C(O)C1NCCC1 | O=C(O)C1[A]CCC1 | **Building block** | [7, 8] |
| 75 | Proline-$CO_2$ | Pyrrolidines | | | N1CCCC1 | [A]1CCCC1 | **Building block** | |
| 76 | Ectoine | Amino acids, peptides, and analogues | Microalgae | *Picochlorum oklahomensis* | CC1=NCCC(C(O)=O)N1 | CC1=NCCC(C(O)=O)[A]1 | **Building block** | [9] |
| 77 | Ectoine-$CO_2$ | Pyrimidines and pyrimidine derivatives | | | CC1=NCCCN1 | CC1=NCCC[A]1 | **Building block** | |
| 78 | Homocysteic acid | Amino acids, peptides, and analogues | Red alga | *Palmaria palmata* | NC(CCS(=O)(O)=O)C(O)=O | [A]C(CCS(=O)(O)=O)C(O)=O | **Building block** | [5] |
| 79 | Homocysteic acid-$CO_2$ | Pyrimidines and pyrimidine derivatives | | | NCCCS(=O)(O)=O | [A]CCCS(=O)(O)=O | **Building block** | |
| 80 | 4-Hydroxyproline | Amino acids, peptides, and analogues | Marine algae | *Cricosphaera carterae*, *Gymnodinium* sp., | OC1CC(C(O)=O)NC1 | OC1CC(C(O)=O)[A]C1 | **Building block** | [10, 11] |

| | | | | | | | | |
|---|---|---|---|---|---|---|---|---|
| | | | | *Skeletonema costatum*, *Nitzschia seriata*, … | | | | |
| 81 | 4-Hydroxyproline-$CO_2$ | Pyrrolidines | | | OC1CCNC1 | OC1CC[A]C1 | **Building block** | |
| 82 | 3,5-Diiodotyrosine | Amino acids, peptides, and analogues | Brown algae | Laminariales | IC1=C(O)C(I)=CC(CC(N)C(O)=O)=C1 | IC1=C(O)C(I)=CC(CC([A])C(O)=O)=C1 | **Building block** | [10, 12] |
| 83 | 3,5-Diiodotyrosine-$CO_2$ | Phenethylamines | | | IC1=C(O)C(I)=CC(CCN)=C1 | IC1=C(O)C(I)=CC(CC[A])=C1 | **Building block** | |
| 84 | 3,5-Diiodothyronine | Amino acids, peptides, and analogues | Brown algae | Laminariales | OC1=CC=C(OC2=C(I)C=C(CC(N)C(O)=O)C=C2I)C=C1 | OC1=CC=C(OC2=C(I)C=C(CC([A])C(O)=O)C=C2I)C=C1 | **Building block** | [10, 12] |
| 85 | 3,5-Diiodothyronine-$CO_2$ | Diphenylethers | | | OC1=CC=C(OC2=C(I)C=C(CCN)C=C2I)C=C1 | OC1=CC=C(OC2=C(I)C=C(CC[A])C=C2I)C=C1 | **Building block** | |
| 86 | Gigartinine | Amino acids, peptides, and analogues | Red algae | Gigartinales: *Gymnogongrus flabelliformis*, *Chondrus crispus*; Gracilariales: *Gracilaria* spp.; Halymeniales: *Polyopes prolifer*, *Grateloupia* spp.; … | O=C(O)C(N)CCCNC(/N=C(N)/N)=O | O=C(O)C([A])CCCNC(N=C(N)N)=O | **Building block** | [5] |
| 87 | Gigartinine-$CO_2$ | Ureas | | | NCCCCNC(/N=C(N)/N)=O | [A]CCCCNC(N=C(N)N)=O | **Building block** | |
| 88 | Lividine | Amino acids, peptides, and analogues | Red algae | *Grateloupia livida*, *Grateloupia filicina* | O=C(O)C(N)CCCNC(NC(N)=O)=O | O=C(O)C([A])CCCNC(NC(N)=O)=O | **Building block** | [5] |
| 89 | Lividine-$CO_2$ | Ureas | | | NCCCCNC(NC(N)=O)=O | [A]CCCCNC(NC(N)=O)=O | **Building block** | |

| | | | | | | | | |
|---|---|---|---|---|---|---|---|---|
| 90 | Carnosine | Hybrid peptides | Red algae, green algae, microalgae | Bangiales: *Neopyropia yezoensis*; Gigartinales: *Sarconema filiforme*, *S. scinaioides*, *Hypnea musciformis*; Ceramiales: *Acanthophora nayadiformis*; *Acrosiphonia saxatilis* | O=C(O)C(NC(CCN)=O)CC1=CN=CN1 | O=C(O)C(NC(CC[A])=O)CC1=CN=CN1 | **Building block** | [5] |
| 91 | Carnosine-$CO_2$ | Imidazoles | | | O=C(NCCC1=CN=CN1)CCN | O=C(NCCC1=CN=CN1)CC[A] | **Building block** | |
| 92 | Anserine | Hybrid peptides | Rare occurrence (one red alga, one brown alga, one green microalga) | *Neopyropia yezoensis*, *Polycladia indica*, *Chlorolobion braunii* | O=C(O)C(NC(CCN)=O)CC1=CN=CN1C | O=C(O)C(NC(CC[A])=O)CC1=CN=CN1C | **Building block** | [5] |
| 93 | Anserine-$CO_2$ | Amino acids, peptides, and analogues | | | O=C(NCCC1=CN=CN1C)CCN | O=C(NCCC1=CN=CN1C)CC[A] | **Building block** | |
| 94 | Ophidine | Hybrid peptides | Marine animals (whales) | | O=C(O)C(NC(CCN)=O)CC1=CN(C=N1)C | O=C(O)C(NC(CC[A])=O)CC1=CN(C=N1)C | **Building block** | [13] |
| 95 | Ophidine-$CO_2$ | Imidazoles | | | O=C(NCCC1=CN(C=N1)C)CCN | O=C(NCCC1=CN(C=N1)C)CC[A] | **Building block** | |
| 96 | Pyrrolidine-2,4-dicarboxylic acid | Amino acids, peptides, and analogues | Red algae | *Chondria coerulescens*, *C. dasyphylla*, *Ceramium virgatum* | O=C(C1NCC(C(O)=O)C1)O | O=C(C1[A]CC(C(O)=O)C1)O | **Building block** | [5] |

| | | | | | | | | |
|---|---|---|---|---|---|---|---|---|
| 97 | Pyrrolidine-2,4-dicarboxylic acid-$CO_2$ | Pyrrolidine carboxylic acids and derivatives | | | O=C(C1CCNC1)O | O=C(C1CC[A]C1)O | **Building block** | |
| 98 | Pipecolic acid | Amino acids, peptides, and analogues | Green microalgae | *Haematococcus lacustris* | O=C(O)C1CCCCN1 | O=C(O)C1CCCC[A]1 | **Building block** | [5] |
| 99 | Pipecolic acid-$CO_2$ | Piperidines | | | C1CNCCC1 | C1C[A]CCC1 | **Building block** | |
| 100 | 5-Hydroxypipecolic acid | Amino acids, peptides, and analogues | Brown algae | *Undaria pinnatifida*, *Pelvetia canaliculata* | O=C(O)C1CCC(O)CN1 | O=C(O)C1CCC(O)C[A]1 | **Building block** | [5] |
| 101 | 5-Hydroxypipecolic acid-$CO_2$ | Piperidines | | | OC1CNCCC1 | OC1C[A]CCC1 | **Building block** | |
| 102 | Baikiain | Amino acids, peptides, and analogues | Red algae | Corallinales, Gigartinales, Ceramiales, Gelidiales, *Gracilaria secundata*, *Sinaia furcellata*, … | O=C(O)C1CC=CCN1 | O=C(O)C1CC=CC[A]1 | **Building block** | [5] |
| 103 | Baikiain-$CO_2$ | Hydropyridines | | | C1CNCC=C1 | C1C[A]CC=C1 | **Building block** | |
| 104 | Azetidine-2-carboxylic acid | Amino acids, peptides, and analogues | Red algae | *Lophocladia trichoclados* | O=C(O)C1NCC1 | O=C(O)C1[A]CC1 | **Building block** | [5] |
| 105 | Azetidine-2-carboxylic acid-$CO_2$ | Azetidines | | | C1CNC1 | C1C[A]C1 | **Building block** | |
| 106 | Kainic acid | Amino acids, peptides, and analogues | Red algae | Ceramiales: *Digenea simplex*, *Palisada perforata*, *Osmundaria obtusiloba*, …; *Palmaria palmata* | CC(C1CNC(C(O)=O)C1CC(O)=O)=C | CC(C1C[A]C(C(O)=O)C1CC(O)=O)=C | **Building block** | [5] |

| | | | | | | | | |
|---|---|---|---|---|---|---|---|---|
| 107 | Kainic acid-$CO_2$ | Pyrrolidines | | | CC(C1CNCC1CC(O)=O)=C | CC(C1C[A]CC1CC(O)=O)=C | **Building block** | |
| 108 | 1´-Hydroxykainic acid | Amino acids, peptides, and analogues | Red algae | *Palmaria palmata* | CC(C)(O)C1CNC(C(O)=O)C1CC(O)=O | CC(C)(O)C1C[A]C(C(O)=O)C1CC(O)=O | **Building block** | [5] |
| 109 | 1´-Hydroxykainic acid-$CO_2$ | Fatty acids and conjugates | | | CC(C)(O)C1CNCC1CC(O)=O | CC(C)(O)C1C[A]CC1CC(O)=O | **Building block** | |
| 110 | Domoic acid | Amino acids, peptides, and analogues | Red algae | *Amansia glomerata*, *Chondria armata*, *C. baileyana*, *Alsidium corallinum* | C/C(C1CNC(C(O)=O)C1CC(O)=O)=C/C=C/C(C)C(O)=O | C/C(C1C[A]C(C(O)=O)C1CC(O)=O)=C/C=C/C(C)C(O)=O | **Building block** | [5] |
| 111 | Domoic acid-$CO_2$ | Fatty acids and conjugates | | | C/C(C1CNCC1CC(O)=O)=C/C=C/C(C)C(O)=O | C/C(C1C[A]CC1CC(O)=O)=C/C=C/C(C)C(O)=O | **Building block** | |
| 112 | 4-Hydroxykainic acid | Amino acids, peptides, and analogues | Red algae | Ceramiales | CC(C1(O)CNC(C(O)=O)C1CC(O)=O)=C | CC(C1(O)C[A]C(C(O)=O)C1CC(O)=O)=C | **Building block** | [5] |
| 113 | 4-Hydroxykainic acid-$CO_2$ | Fatty acids and conjugates | | | CC(C1(O)CNCC1CC(O)=O)=C | CC(C1(O)C[A]CC1CC(O)=O)=C | **Building block** | |
| 114 | Laminine | Amino acids, peptides, and analogues | Brown algae, less frequently in red algae | Laminariales: *Laminaria*, *Kjellmaniella*, *Undaria*; Fucales: *Fucus* spp., *Ascophyllum nodosum*, …; *Ahnfeltia* spp., *Gymnogongrus griffithsiae*, … | NC(CCCC[N+](C)(C)C)C(O)=O | [A]C(CCCC[N+](C)(C)C)C(O)=O | **Building block** | [5] |
| 115 | Laminine-$CO_2$ | Quaternary ammonium salts | | | NCCCCC[N+](C)(C)C | [A]CCCCC[N+](C)(C)C | **Building block** | |

| | | | | | | | | |
|---|---|---|---|---|---|---|---|---|
| 116 | Lanthionine | Amino acids, peptides, and analogues | Red algae | Gigartinales: *Hypnea musciformis* | NC(CSCC(C(O)=O)N)C(O)=O | NC(CSCC(C(O)=O)[A])C(O)=O | **Building block** | |
| 117 | Lanthionine-$CO_2$ | Amino acids, peptides, and analogues | | | NC(CSCCN)C(O)=O | NC(CSCC[A])C(O)=O | **Building block** | |
| 118 | Rhodoic acid (tauropine) | Amino acids, peptides, and analogues | Red algae | Gigartinales: *Chondrus* spp., *Mazzaella* spp., *Neodilsea yendoana*, …; Ceramiales: *Laurencia glandulifera*, *Neorhodomela larix*, …; Bangiales: *Pyropia* spp.; Gelidiales: *Gelidium amansii* | CC(NCCS(O)(=O)=O)C(O)=O | CC([A]CCS(O)(=O)=O)C(O)=O | **Building block** | [5] |
| 119 | Rhodoic acid-$CO_2$ | Organosulfonic acids and derivatives | | | CCNCCS(O)(=O)=O | CC[A]CCS(O)(=O)=O | **Building block** | |
| 120 | Methionine sulfoxide | Amino acids, peptides, and analogues | Brown alga, green algae, red algae | Ectocarpales, Fucales, Laminariales, Ulvales, Cladophorales, Gigartinales, Ceramiales, Halymeniales, … | NC(CCS(C)=O)C(O)=O | [A]C(CCS(C)=O)C(O)=O | **Building block** | [5] |
| 121 | Methionine sulfoxide-$CO_2$ | Sulfoxides | | | NCCCS(C)=O | [A]CCCS(C)=O | **Building block** | |

| | | | | | | | | |
|---|---|---|---|---|---|---|---|---|
| 122 | Chondrine (yunaine) | Amino acids, peptides, and analogues | Brown alga, green algae, red algae | *Chondria crassicaulis* (first organism where it was discovered), Ceramiales, Gigartinales, Laminariales, Fucales, … | O=S1CC(C(O)=O)NCC1 | O=S1CC(C(O)=O)[A]CC1 | **Building block** | [5] |
| 123 | Chondrine-$CO_2$ | Thiomorpholines | | | O=S1CCNCC1 | O=S1CC[A]CC1 | **Building block** | |
| 124 | Cysteinolic acid | Organosulfonic acids and derivatives | Microalgae, brown algae, green algae, red algae, … | *Thalassiosira weissflogii*, *Chondria crassicaulis*, *Sargassum serratifolium*, *Vertebrata lanosa*, … | O=S(CC(N)CO)(O)=O | O=S(CC([A])CO)(O)=O | **Building block** | [14, 15] |
| 125 | β-Glutamate | Fatty acids and conjugates | Archaea | *Methanothermococcus thermolithotrophicus*, … | NC(CC(O)=O)CC(O)=O | [A]C(CC(O)=O)CC(O)=O | **Building block** | [16] |
| 126 | β-Glutamate-$CO_2$ | Amino acids, peptides, and analogues | | | NC(CC(O)=O)C | [A]C(CC(O)=O)C | **Building block** | |
| 127 | Hydroxyectoine | Amino acids, peptides, and analogues | Bacteria | *Halomonas elongata*, *Nocardiopsis halophila* | O=C(O)C1N=C(NCC1O)C | O=C(O)C1N=C([A]CC1O)C | **Building block** | [16] |
| 128 | Hydroxyectoine-$CO_2$ | Pyrimidines and pyrimidine derivatives | | | OC1CN=C(NC1)C | OC1CN=C([A]C1)C | **Building block** | |

| | | | | | | | | |
|---|---|---|---|---|---|---|---|---|
| 129 | Cysteamide | Amino acids, peptides, and analogues | Squid, oyster | *Loligo pealii*, *Dosidicus gigas*, *Ostrea edulis* | O=C(N)C(CS(=O)(O)=O)N | O=C(N)C(CS(=O)(O)=O)[A] | **Building block** | [15] |
| 130 | Cysteic acid | Amino acids, peptides, and analogues | Plants, fungi | (precursor of cysteinolic acid) | O=C(O)C(CS(=O)(O)=O)N | O=C(O)C(CS(=O)(O)=O)[A] | **Building block** | [15] |
| 131 | Cysteic acid-$CO_2$ | Organosulfonic acids and derivatives | | | NCCS(=O)(O)=O | [A]CCS(=O)(O)=O | **Building block** | |
| 132 | AHPS (Aminohydroxy-propanesulfonate) | Organosulfonic acids and derivatives | Red algae | *Grateloupia livida* | NCC(O)CS(O)(=O)=O | [A]CC(O)CS(O)(=O)=O | **Building block** | [15] |
| 133 | Homotaurine | Organosulfonic acids and derivatives | Brown algae, red algae, green algae | *Fucus vesiculosus*, *Ascophyllum nodosum*, *Chondrus crispus*, *Gracilaria longissima*, *Palmaria palmata*, … | NCCCS(=O)(O)=O | [A]CCCS(=O)(O)=O | **Building block** | [15, 17] |
| 134 | Nγ-Acetyldiaminobutyric acid | Amino acids, peptides, and analogues | Bacteria | *Halomonas elongata* | CC(NCCC(C(O)=O)N)=O | CC(NCCC(C(O)=O)[A])=O | **Building block** | [16] |
| 135 | Nγ-Acetyldiaminobutyric acid-$CO_2$ | Carboxylic acid derivatives | | | CC(NCCCN)=O | CC(NCCC[A])=O | **Building block** | |
| 136 | Nε-Acetyl-β-lysine | Amino acids, peptides, and analogues | Archaea | *Methanothermococcus thermolithotrophicus*, … | CC(NCCCC(N)CC(O)=O)=O | CC(NCCCC([A])CC(O)=O)=O | **Building block** | [16] |
| 137 | Nε-Acetyl-β-lysine-$CO_2$ | Carboxylic acid derivatives | | | CC(NCCCC(N)C)=O | CC(NCCCC([A])C)=O | **Building block** | |

### 12.Literature
